# From injury to recovery: functional neuronal regeneration after traumatic brain injury in the telencephalon of the young adult killifish

**DOI:** 10.1101/2025.08.28.672347

**Authors:** Valerie Mariën, Caroline Zandecki, Jolien Van houcke, Anouk Maes, Rajagopal Ayana, Chris Van den Haute, Rik Gijsbers, Marialuisa Tognolina, Lutgarde Arckens

## Abstract

Neuronal loss caused by neurodegenerative diseases and traumatic brain injuries (TBI) often results in long-term disabilities, highlighting the urgent need for further research and effective regenerative strategies. In mammals, neurogenic capacity is inherently limited and declines further with age. In contrast, the young adult killifish demonstrates a remarkable ability to regenerate neurons in the telencephalon following TBI. However, it remains unknown whether and when these newly generated neurons functionally integrate into existing circuits, as traditional histological analysis of fixed tissue offers only static insights into this dynamic process. To this end, we optimized a retroviral vector strategy to label dividing stem cells and their progeny, including newborn neurons. By introducing a combination of novel approaches i.e., retroviral vector labeling, electrophysiology and a conditioned place avoidance test, we investigated the generation, morphology, and synaptic integration of newborn neurons following TBI in the dorsomedial (Dm) zone of the telencephalon, a region homologous to the mammalian amygdala in other teleost fish. Our results show that injury-induced adult-born neurons functionally integrate into existing circuits, and that killifish can achieve functional behavioral recovery after TBI. While previous histological assessments using a stab-wound injury suggested a 30-day recovery period, our functional data reveal that full behavioral recovery requires approximately 50 days. At this point, fish successfully relearn to avoid a conditioned place, and the new neurons exhibit mature morpho-electric characteristics, including abundant dendritic spines. Electrophysiological analysis revealed that newborn neurons in an injured environment take longer to mature when compared to neurons in naive killifish. Together, our findings demonstrate that structural regeneration aligns with functional recovery, and establish retroviral vectors as a powerful tool for birth dating injury-induced neurogenesis in teleosts. Killifish thus represent a promising model for studying interventions aimed at enhancing neuronal maturation and integration after brain injury.

**Key points:**

- A retroviral vector strategy allows specific and sparse labeling of adult-born neurons in the killifish brain.
- The Dm zone in the telencephalon of the killifish is responsible for avoidance learning and memory and thus homologous to the mammalian amygdala.
- In young adult killifish, upon Dm injury, adult-born neurons mature morphologically and functionally in 50 days, which is slower than in constitutive neurogenesis.
- The timing and extent of behavioral recovery from such injury aligns with morpho-electric observations.

## 1 Introduction

Neuronal loss in the context of neurodegenerative diseases and traumatic brain injuries often results in long term disabilities. Restoring brain function upon neuronal loss due to pathology or trauma requires the induction of a high level of neuroplasticity through *de novo* neurogenesis, cell migration, differentiation, and circuit integration, which involves neurite outgrowth and synaptogenesis. Unfortunately, the mammalian brain has a very limited neurogenic capacity [1], that is restricted to two neurogenic niches; the subventricular zone of the lateral ventricles and the subgranular zone of the dentate gyrus in the hippocampus [2–5]. Nevertheless, the mammalian brain is capable of adult neurogenesis upon injury [6,7]. Stem cells divide and generate newborn neurons, but these are often not able to properly mature and integrate into the pre-existing network, which hampers functional recovery after brain injury [6–8]. In addition, a glial scar is formed upon injury, restricting neurons from migrating towards the injury site and hence limiting functional recovery [9].

Teleost fish have emerged as valuable models in regenerative biology due to their remarkable ability to repair various organs, including the brain, throughout their life. This regenerative capacity is likely linked to their indeterminate growth and the sustained activity of neural stem cell niches, in sharp contrast to the determinate body growth and limited neurogenesis observed in humans [10,11]. Notably, the African turquoise killifish (*Nothobranchius furzeri*), contrary to other teleostean model species such as zebrafish, only shows high potency to regenerate at a young age. While young killifish demonstrate robust recovery from traumatic brain injury (TBI), aged individuals exhibit a marked decline in regenerative capacity, including the formation of persistent glial scarring, mirroring the mammalian condition [12]. This unique teleost model thus holds great promise in revealing new knowledge on how potent regeneration is diminished upon aging and on how this might be restored.

Our previous findings suggest that potent recovery in young killifish may be driven by the generation of a substantially larger pool of newborn neurons, as compared to aged fish, that are capable of migrating into the brain parenchyma [12]. However, whether such newborn neurons truly become functionally integrated into the existing circuitry to drive normal behavior, even at a young age, remained unexplored due to the lack of available tools applicable to killifish. In this study, we optimized a triple approach to address this gap. To assess the timing and extent of morphological and functional maturation of injury-induced adult-born neurons in the killifish telencephalon, we first optimized a retroviral vector strategy, based on a previous report in zebrafish, to selectively label dividing progenitor cells and their progeny via GFP expression [13]. We then combined this with patch-clamp electrophysiology in *ex vivo* brain explants to record from GFP+ adult-born neurons. Finally, we implemented a behavioral assay of functional recovery using a conditioned place avoidance test. Successful performance in this learning and memory test requires an intact and functional amygdala, which we presumed to correspond to the killifish dorsomedial (Dm) zone based on knowledge from other teleosts [14].

Here, we show that adult killifish can functionally regenerate after TBI by 50 dpi and that by then injury-induced newborn neurons morpho-electrically resemble newborn neurons in naive killifish. These newborn neurons receive synaptic input, showing integration into the neural circuitry. Moreover, we provide behavioral evidence that the killifish Dm zone is involved in fear learning and relearning upon Dm injury and hence homologous to the mammalian amygdala similar to other teleost fish. Our optimized research pipeline holds great promise for the discovery of new regenerative strategies in killifish with future relevance to human brain repair.

## 2 Results

### 2.1 A retroviral vector strategy allows birth dating of adult-born neurons following a slit-injury in the killifish telencephalon

To expedite investigating newborn neuron maturation and integration, we first established a reproducible slit-injury model in the killifish telencephalon. The Dm zone was chosen as target area since this brain region is implicated in fear learning and memory in other teleost fish. To verify the reproducibility of the slit-injury, we measured the injury depth and the distance from the midline and compared this to the killifish brain atlas [15]. In ±70% of the analyzed sections, the injury was inflicted to the target area (n= 5 fish, n=20 sections), with a mean depth of 244 µm (minimal-maximal injury depth of 106 to 466 µm) and a mean distance from the midline of 178 µm (minimal-maximal distance to the midline of 130 µm to 246 µm) (Supplementary Fig. 1).

Next, we performed a series of tests to confirm that the γ-retroviral (RV) MLV vector (MLV-EFs-eGFP) effectively labels dividing progenitors and their progeny in the killifish telencephalon following injection into the telencephalic ventricle (Fig. 1A, B). To confirm that this type of vector only integrates in dividing cells in killifish, like in mammals, we combined viral vector transduction with a 4-day EdU pulse labeling. We quantified the degree of double labeling in naive and injured fish (Supplementary Fig. 2), which amounted to 70-90%. In the remaining GFP+ cells (10-30%), EdU was not detectable (Supplementary Fig. 2L), and a clear variability in EdU labeling intensity was observed among the different GFP+ cells (Supplementary Fig. 2E, I). Given the 21-day chase period, during which multiple rounds of cell division may have occurred, it is likely that the EdU signal became progressively diluted [16]. Notably, all GFP+/EdU-cells were located near the ventricular zone, surrounded by EdU+ cells, and exhibited morphologies similar to those of the double-positive population (Supplementary Fig. 3). Taken together, these observations strongly support the conclusion that the retroviral vector was incorporated exclusively in dividing progenitors and passed on to their progeny. The limited number of GFP+/EdU-cells can be attributed to successive cell divisions that diluted the EdU signal below the detection threshold. In sum, the MLV vector approach provides a reliable means of labeling dividing progenitor cells and their progeny in the adult killifish telencephalon.

**Figure 1:**
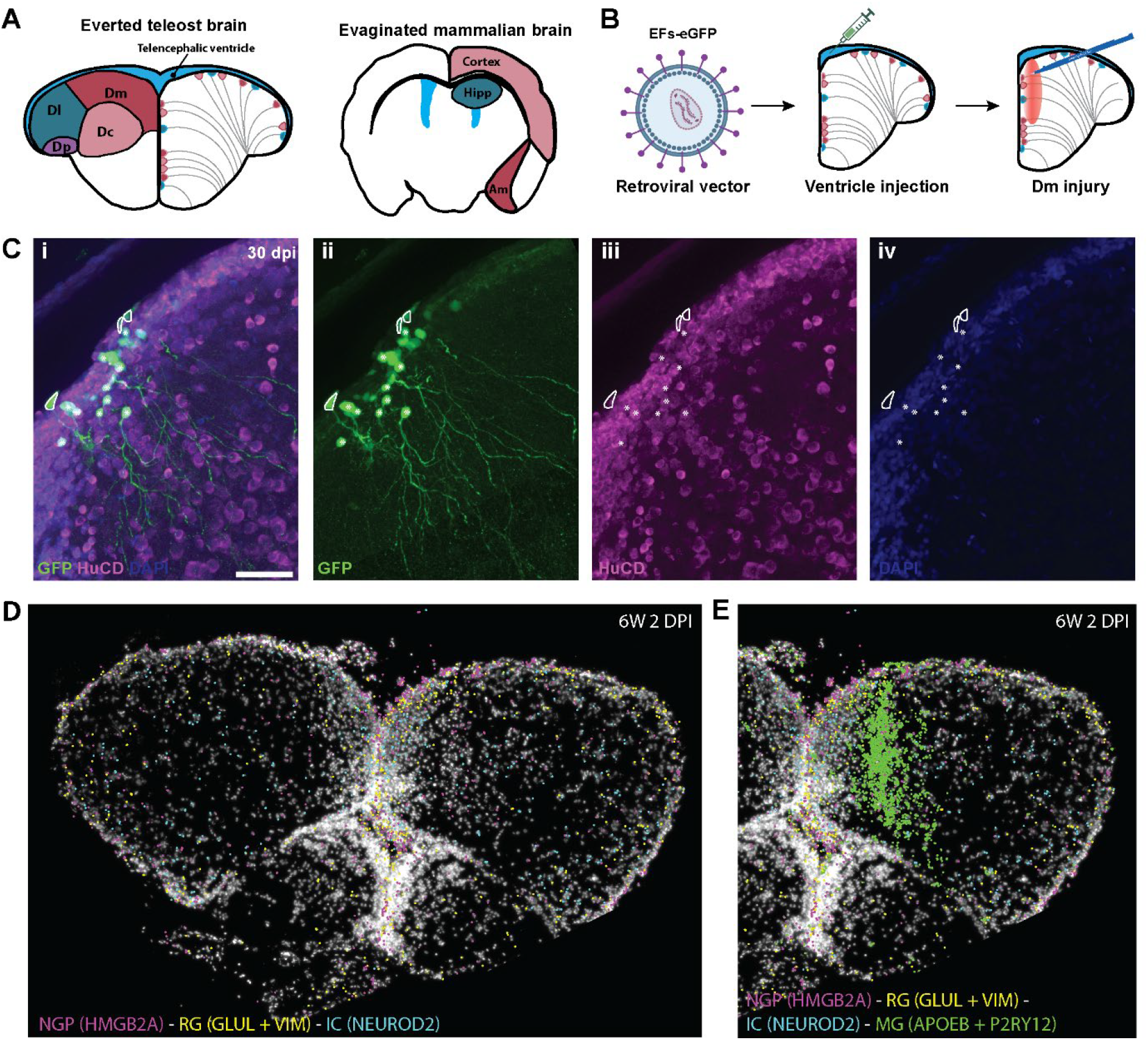
Retroviral vectors as a tool to study the generation and maturation of newborn neurons, generated from dividing NGPs as verified by in situ sequencing. (A) The teleost brain is everted during development, which results in the orientation of the ventricle on the outside of the telencephalon as well as the stem cells lining the telencephalic ventricle. The homologous regions between the teleost and the mammalian brain are color-coded on the coronal schematics. (B) In mammals, RV vectors only integrate into dividing cells. A RV vector expressing eGFP under control of universal human EF1a short promoter (EFs) was injected into the telencephalic ventricle. Upon injection in killifish, dividing stem cells are predicted to also incorporate the viral vector construct into their DNA. Upon further proliferation, differentiation and maturation, the newborn neurons will then be fluorescently labeled (GFP+). Next, an injury was given into the Dm zone from rostral to caudal with a micro knife. (C) Co-labeling for HuC/D confirmed the neuronal nature of the GFP+ cells, here at 30 days post-injury. Cells marked with an asterisk are adult-born neurons (GFP+ HuC/D+), cells outlined are stem cells, recognizable by their triangular cell body, and the location at the ventricular zone. Scale bar: C(i): 200 µm; C(ii, iii, iv): 10 µm; E: 50 µm. RV: retroviral. (D) In situ sequencing of the telencephalon at 2 dpi reveals an increase in (dividing) NGPs, RG and IC medial to the Dm injury. (E) Addition of the microglia/macrophage (MG/MF) markers APOEB and P2RY12 reveals the location of the injury and shows a strong inflammatory response. NGPs: non-glial progenitors, RG: radial glia, IC: intercell.

To determine whether the GFP+ cells mature into neurons, we performed immunohistochemical double stainings for GFP and HuC/D (Elavl3/4), a pan-neuronal marker that labels neuroblasts, immature and mature neurons [17]. As expected, the majority of GFP+ cells expressed HuC/D, indicating neuronal identity (Fig. 1C). Using intraventricular retroviral vector injections, we can thus study and birthdate newborn neurons in the killifish telencephalon.

We also observed GFP+ cells lacking HuC/D expression, which were lining the ventricular surface and hence are likely the founder stem cells. Their triangular cell body (outlined in white; Fig. 1C) is typically observed for progenitor/glial cells [10,12,18]. In killifish, adult-born neurons are derived from the highly-proliferative NGPs, and not from radial glia like in zebrafish, as demonstrated in our previous work [12,17] and by others [18]. Another subset of GFP+/HuC/D- cells were located close to the ventricle and had rounder cell bodies, and are most likely early differentiating progenitor cells.

We performed in situ sequencing analysis to visualize where alterations in cell division occur, as well as in the abundance of progenitor cells and a type of intercell, intercell (IC)-neuronal cell (NC) [17]. These IC-NCs were previously predicted, through lineage inference, to develop from NGPs and to differentiate into immature and mature neurons in the young adult killifish (Fig. 1D, Supplementary Fig. 4). Comparison of 6w naive and 6w-2dpi fish revealed increased expression of the division markers PCNA and MKI67 in HMGB2A-positive NGPs at 2 dpi, with the highest expression level in the right lesioned hemisphere of 2 dpi fish at the ventricular surface of the Dm zone (Supplementary Fig. 4A). Expression in the intercell-NC marker gene NEUROD2 was clearly increased just below this region (Fig. 1D, Supplementary Fig. 4F), indicating the rapid formation of a substantial amount of Intercell-NC cells already at 2 dpi. Increased GAP43 expression at 2 dpi further substantiates localized intensified neurogenesis in the lesioned hemisphere very early upon injury (Supplementary Fig. 4C, F). APOEB and P2RY12, marker genes for microglia/macrophages (MG/MF), reveal MG/MF proliferation and activation at the injury site at 2 dpi [12,17], as well as within the nearby region of neuroregeneration (Supplementary Fig. 4D, F). In addition to MG/MF, the increase in VIM- and GLUL-positive astroglia (RG), interspersed with NGPs, also reflects local glial proliferation/activation specifically in the lesioned Dm zone (Fig. 1D, Supplementary Fig. 4E, F). Together these molecular observations of distinct cellular responses medial from the Dm injury site guided our follow-up experiments, enabling the use of a viral vector-based sparse labeling strategy to investigate the morpho-electric maturation of injury-induced neurons born in the neurogenic niche closest to the injury site and relevant to the creation of new excitatory neurons for the Dm region of the telencephalon [10].

### 2.2 Adult-born neurons undergo morphological maturation upon injury

To study the morphological properties of newborn neurons post-injury, and compare them with those in naive fish, two different viral vectors were used for targeted labeling. First, in 6-week old naive killifish, an AAV2.1 vector (CAG-eGFP) was used and injected directly and deep into the brain parenchyma, targeting regions where mature neurons reside [10]. Unlike the MLV RV vector, AAV2.1 can transduce both dividing and non-dividing cells and is therefore well-suited for labeling fully mature neurons [10]. The injection strategy takes advantage of the telencephalon’s concentric growth pattern, enabling selective labeling of older, centrally located neurons, which are typically born within the first two weeks after hatching [10]. Second, in injured killifish, the MLV (EFs-eGFP) RV vector was injected into the telencephalic ventricle to selectively and sparsely label dividing progenitor cells/NGPs lining the ventricular surface. This approach allowed us to track the morphological maturation of their progeny at multiple time points post-injury: 11, 18, 30, 36, 50, and 70 dpi (Fig. 2A). The morphology of these labeled neurons was then compared to that of mature Dm neurons in naive fish labeled with the AAV.1 vector.

**Figure 2:**
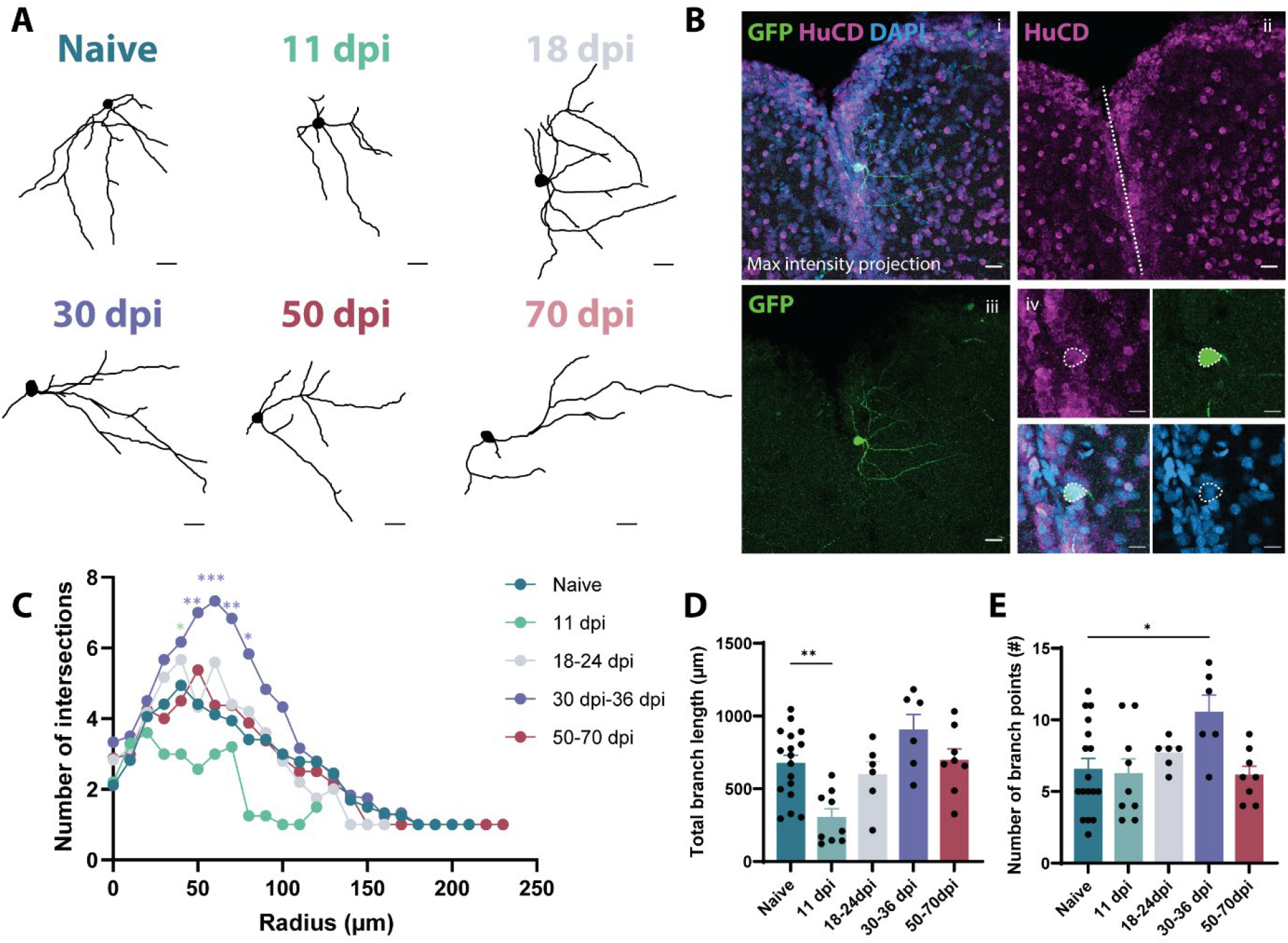
By 50 dpi, newborn neurons in injured fish exhibit morphologies comparable to those of mature neurons in naive fish. (A) Skeletons of confocal-traced Dm neurons in a naive fish and at different post-injury timepoints. (B) (i,ii,iii) Maximal intensity projection of an 18 dpi RV-vector labeled neuron, scale: 20 µm. GFP co-stained with HuC/D (neuronal marker) and DAPI (nuclei). (iv) Single confocal plane showing double labeling for GFP and HuC/D. Scale: 10 µm. (C) Sholl graph for the reconstructed cells per condition. At 11 dpi a difference in the number of intersections was observed with the naive neurons at a radius of 40 µm. For the 30-36 dpi timepoint, a difference was observed with the naive neurons at a radius of 50, 60, 70, 80 µm (Two-way ANOVA, mean plotted). (D) The total branch length was significantly lower at 11 dpi compared to the naive neurons (One-way ANOVA, mean with SEM). (E) The number of branch points was significantly higher at 30-36 dpi compared to neurons in naive fish (One-way ANOVA, mean with SEM). C, D, E: N_naive_= 17, N_11dpi_= 10, N_18-24dpi_= 6, N_30-36dpi_= 6, N_50-70dpi_= 8. Dpi: days post-injury. RV: retroviral.

Sholl analysis of traced neurons revealed that at 11 dpi, newborn neurons of injured fish exhibited significantly shorter neurites compared to mature neurons in naive fish (Fig. 2C, D, p=0.0014, One-way ANOVA, F (4, 40) =6.980). At 30-36 dpi, the number of branch points was higher than in the mature neurons of naive fish (Fig. 2E, p=0.0191, One-way ANOVA, F (4, 40) = 2.853), suggesting a transient phase of exuberant outgrowth. By 50 dpi, branch complexity had returned to levels comparable to neurons in naive fish, indicating the occurrence of a pruning phase. Together, these findings suggest that newborn neurons undergo a period of excessive branching followed by refinement. All traced neuron morphologies are shown in Supplementary Fig. 5.

To determine whether and when the injury-induced newborn neurons would receive presynaptic input, an immunohistochemical staining for the presynaptic marker SV2 was carried out on MLV transduced brain slices (Supplementary Fig. 5). Already at 23 dpi, the newborn neurons in the Dm zone were surrounded by SV2+ synaptic contacts around both their cell body and neurites. We interpreted this as a first indication that by 23 dpi the newborn neurons are being integrated into the circuitry.

### 2.3 Timing of functional maturation of adult-born neurons following injury

#### 2.3.1 Assessing electrophysiological properties of adult-born neurons in naive and injured fish without birth dating information

To study the functional maturation of newborn neurons following injury, we chose to introduce *ex vivo* whole-cell patch clamp recordings as approach. To do so, electrophysiological protocols were optimized for killifish (Fig. 3A-B), and recorded cells were biocytin filled during patching to allow post hoc morphological analysis (Fig. 3C-D). We recorded from cells between 35 and 72 dpi and compared their properties to age-matched naive fish (35-72 dpt, days post-transduction), without accounting for the birth date of the recorded cells (GFP-cells). When we compared cells in different electrophysiology protocols (Fig. 3E-G), no significant differences were observed, except for the average EPSC amplitude, which was lower for the neurons in the injury condition (p=0.0498, two-tailed unpaired t-test, t=2.021, df=41). Additionally, the cells in the injured group exhibited greater heterogeneity and a wider distribution of measured electrophysiological properties compared to those from the naive condition, especially with respect to input resistance. The cells in the injured fish had more characteristics of immature cells, i.e., high input resistance (p=0.0843, Mann Whitney), more depolarized resting membrane potential and low spike amplitude (Fig. 3H-K). These results prompted us to consider correlating the birth date of neurons to the electrophysiological parameters, and hence recording from GFP+ cells.

**Figure 3:**
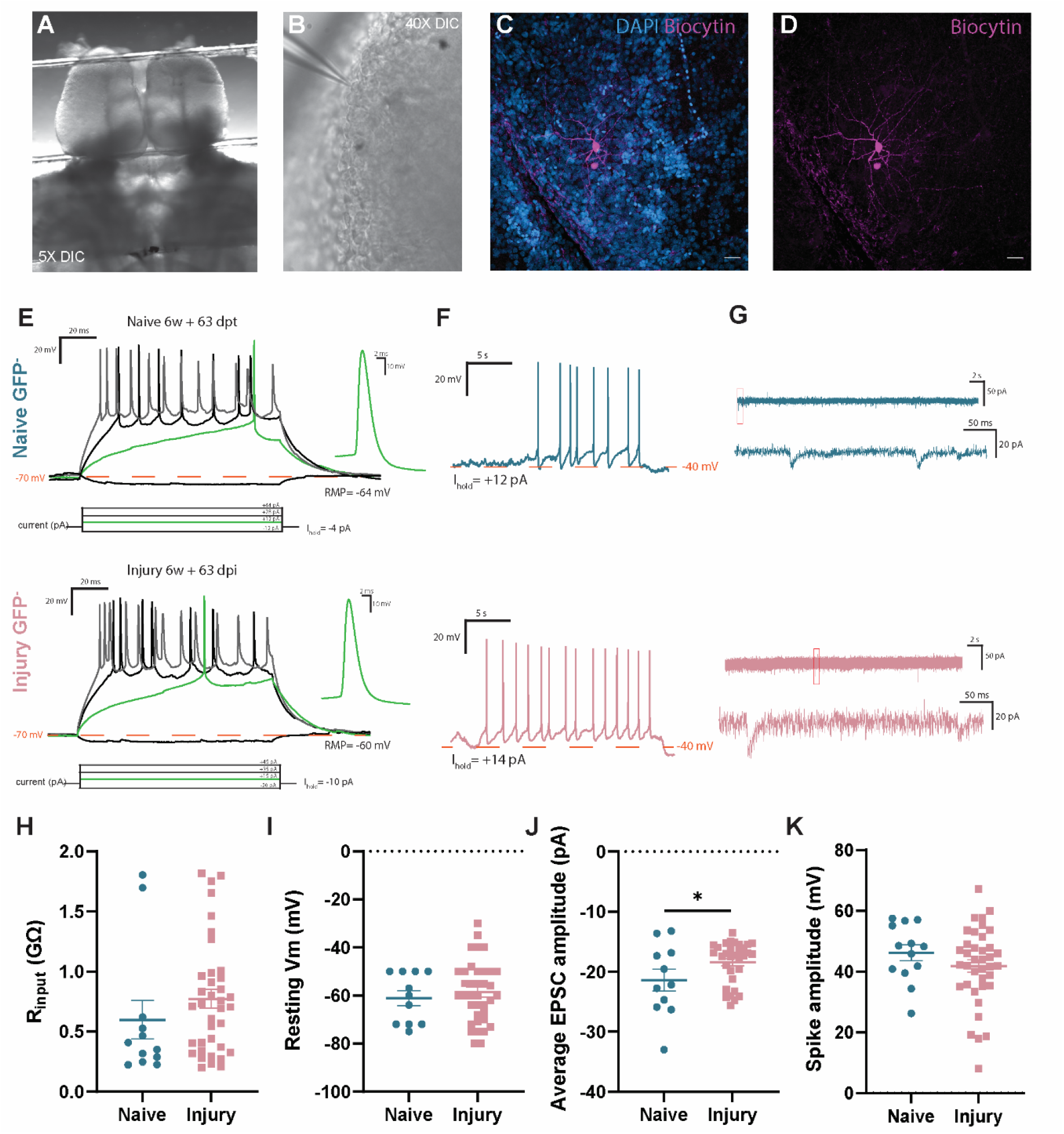
Electrophysiological analysis of neurons after injury shows maturation differences between neurons from naive and injured fish when the birth date is not taken into account (GFP-cells). (A) A killifish *ex vivo* brain explant in artificial cerebrospinal fluid (ACSF) under differential interference contrast (DIC) lighting. (B) The brain in the electrophysiology setup with continuous ACSF perfusion with a 40X objective and DIC light. The patch pipette is visible in the left upper corner. (C-D) During whole cell patch-clamp, the cells are filled with biocytin. After fixation and staining, confocal images can be made of the cell in the full brain. (E) Current clamp (CC) step protocol during whole-cell patch-clamp of a cell at 63 dpt in a naive fish brain (upper row) and at 63 dpi in an injured brain (lower row). Representative membrane potential to hyperpolarizing (bottom black) and depolarizing (green, top black, grey) current injections. Right: first action potential to minimal current injection. (F) Gap-free recording after bringing the membrane potential to a threshold where firing is observed (around -40 mV) in the CC step protocol. (G) Gap-free voltage clamp (VC) protocol at -70 mV recording spontaneous postsynaptic currents. The lower trace is the enlarged view of the boxed area of the top trace. (H-K) Comparison of electrophysiological properties for neurons in naive and post-injury fish brains shows more immature cells in the injury condition as observed by a higher input resistance (p=0.0843, Mann Whitney test), and a larger heterogeneity and wider distribution in the resting membrane potential and spike amplitude. (H) Input resistance, (I) Resting membrane potential, J) Average excitatory postsynaptic current (EPSC) amplitude at -70 mV, (K) Spike amplitude. (H-K) The mean value and standard error of the mean (SEM) is indicated and a significant difference was observed for the average EPSC amplitude (p=0.0498, two-tailed unpaired t-test). N; naive: 12, injury: 37. Dpi: days post-injury, dpt: days post-transduction.

#### 2.3.2 Comparison of electrophysiological parameters in newborn neurons of comparable birth date in naive and injured killifish

To control for the birth date of the cells in naive and injured fish, we selectively recorded only from the sparsely labeled GFP+ cells between 30 and 72 dpi/days post-transduction (dpt) transduced with the MLV approach via ventricular injections. These cells were clearly visible during patch-clamp recordings without the need for immunohistochemical staining in the intact, dissected killifish brains (Fig. 4A). The GFP+ neurons are born from progenitor cells/NGPs dividing at the time of viral vector injection, and hence have a similar birth date in naive and injured fish. After opening the cell and reaching the whole-cell configuration, the GFP in the cell soma would decrease in intensity over time because of the washout with the intracellular solution of the pipette. After 25 minutes in whole-cell configuration, the GFP signal became nearly invisible (Fig. 4B). This served as an additional validation during patch-clamp recordings to ensure that only GFP+ cells were included in the dataset. Following each recording, the pipette was carefully removed and the brain was fixed overnight and subsequently double stained for GFP and biocytin. Morphological reconstructions using confocal imaging revealed the complete morphology of the patched cells and confirmed their newborn identity (GFP+), detectable with anti-GFP staining (Fig. 4C, Fig. 5A). The same electrophysiological protocols were carried out as before (Fig. 4E-G) and, for analysis, cells from naive and injured fish were grouped between 30-50 and 51-72 dpt/dpi. These time windows were based on the outcome of the Sholl analysis (Fig. 2) and the behavioral experiments (Fig. 6). When comparing input resistance, we observed a trend toward a lower input resistance at the later timepoint as compared to the earlier timepoint in both naive and injured fish, suggesting the presence of more mature neurons at the later stages (Fig. 4H). The resting membrane potential did not differ between groups (Fig. 4I). The average amplitude of excitatory postsynaptic currents (EPSCs) was significantly lower at the later post-injury timepoint compared to the earlier timepoint (p=0.0359, Two-way ANOVA, F (1,25) = 4.598), with the later timepoint showing EPSC amplitudes more comparable to those in the naive condition. The maximal spike amplitude and frequency of firing was similar across all conditions (Fig. 4K and data not shown). It thus seems that the injury-induced newborn neurons mature slower compared to naive newborn neurons.

**Figure 4:**
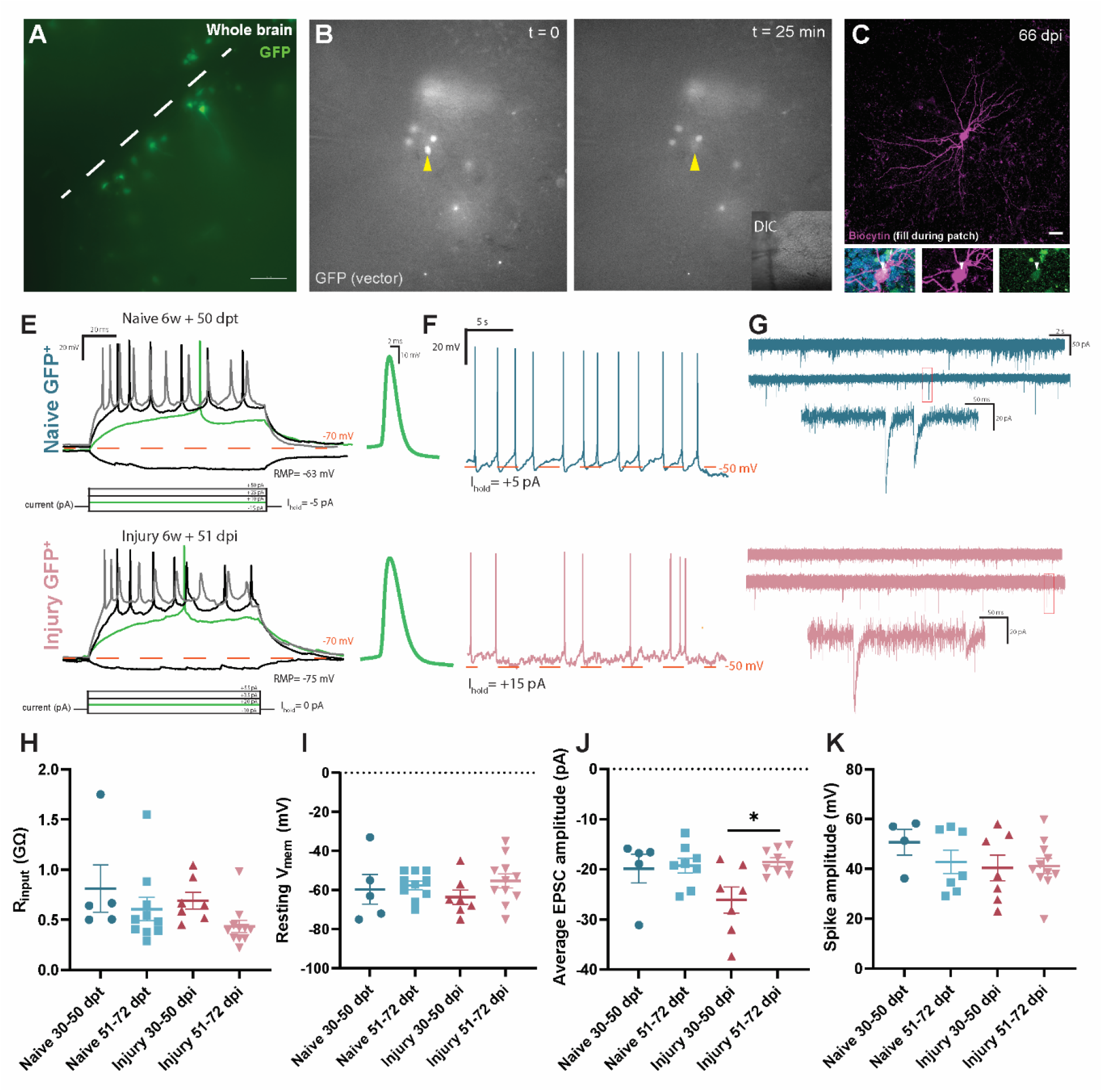
Electrophysiological analysis of adult-born GFP+ neurons reveals a 50-day maturation phase upon injury. (A) A killifish *ex vivo* brain explant showing GFP+ cells under fluorescent light (excitation 395 nm). White dotted line indicates the midline between the two telencephalic hemispheres. (B) GFP+ cells lose their GFP signal upon patching because of the dilution with the intracellular solution. After 25 min, the GFP signal is no longer visible. Right lower corner shows the location (right Dm zone) of this cell under DIC lighting. (C) During whole cell patch-clamp, the cells are filled with biocytin. After fixation and staining, confocal images can be made of the cell in the full brain showing all neurites and the GFP+ signal in the nucleus. (E) Current clamp (CC) step protocol during whole-cell patch-clamp of a 50 dpt cell in a naive fish brain and a 51 dpi cell in an injured fish brain. Representative membrane potential to hyperpolarizing (bottom black) and depolarizing (green, top black, grey) current injections. Right: first action potential to minimal current injection. (F) Gap-free recording after bringing the membrane potential to a threshold where firing is observed (around -40/-50 mV) in the CC step protocol. (G) Gap-free voltage clamp (VC) protocol at -70 mV recording spontaneous postsynaptic currents. The lower trace is the enlarged view of the boxed area of the top trace. Comparison in electrophysiological properties for adult-born GFP+ neurons in naive fish brains and post-injury during two time windows: 30-50 dpi/dpt and 51-72 dpi/dpt; (H) Input resistance, (I) Resting membrane potential, J) Average excitatory postsynaptic current (EPSC) amplitude at -70 mV, (K) Maximal spike amplitude. (H-K) The mean value with SEM is indicated and a significant difference was observed for the EPSC amplitude between the two injury timepoints (p=0.0359, two-way ANOVA). N; naive_30-50dpt_: 5, naive_51-72dpt_: 7, injury_30-50dpi_: 7, injury_51-72dpi_: 9.

**Figure 5:**
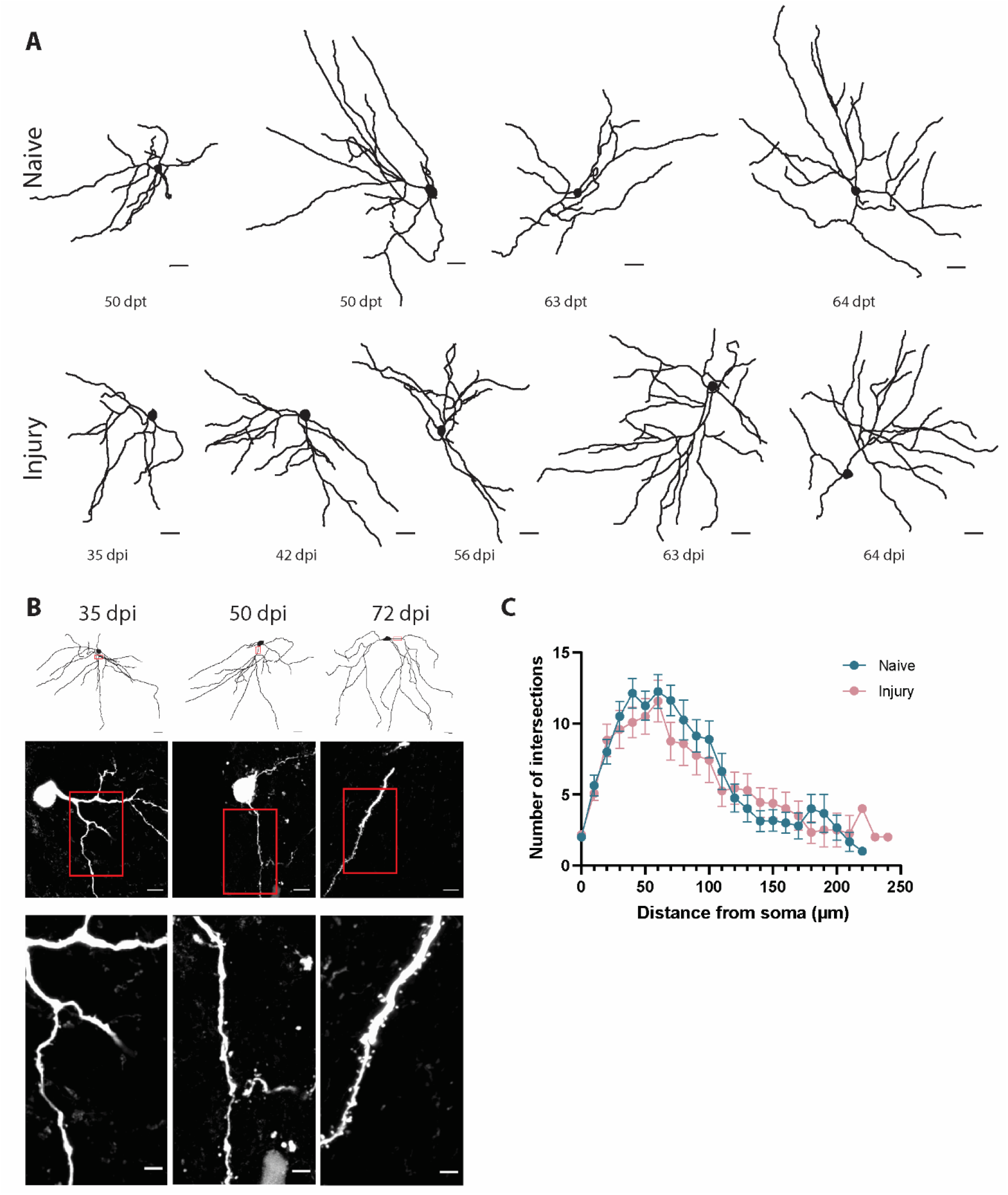
Morphological maturation of adult-born neurons is comparable in the naive and injured brain. Neurons were biocytin filled during patch-clamp electrophysiology. (A) Three-dimensional reconstructions of biocytin filled neurons at different time-points post transduction of the viral vector in fish with (dpi) or without injury (naive, dpt). (B) Spines are present on the neurites and increase gradually from 35 to 72 dpi. Top row shows the 3D skeletons after imaging the full brain. Middle row shows 63X images after coronal sectioning. Bottom row shows an enlarged view of the boxed area in the middle row. Scale bar A: 20 µm, B_top_: 20 µm, B_middle_: 5 µm, B_lower_: 2 µm. (C) Sholl analysis of neurons in naive and injured killifish shows no differences between the two conditions for all timepoints combined (Two-way ANOVA, mean with SEM). C: N_naive_=8, N_injury_=12. Dpi: days post-injury, dpt: days post-transduction.

### 2.4 Morphological maturation and progressive spine formation of newborn neurons following injury

Sholl analysis was conducted on traced biocytin-filled, GFP+ neurons to compare the morphological maturation of neurons in naive versus injured fish (Fig. 5A). Overall, neurons in both conditions exhibited similar morphological features (Fig. 5C). Regardless of whether they were generated during constitutive growth or in response to injury, neurons reached comparable neurite lengths and branching complexity. Notably, the absolute number of detectable branches was higher when analyzing whole brains (up to 17) compared to brain slices (up to 10), highlighting the advantage of preserving the full neuronal architecture. In addition, dendritic spines began to appear on injury-induced neurons around 42 dpi, with only sparse spines observed prior to this time point (Fig. 5B). At later stages, around 66 and 70 dpi, a marked increase in spine density was observed. In sum, these data combined with the data created from sections (Fig. 2) indicate that injury-induced newborn neurons achieve full morphological maturity, including complex branching and spine formation, by approximately 50 dpi, following an overgrowth phase around 30-36 dpi.

### 2.5 Functional behavioral recovery aligns with morpho-electric maturation of adult-born neurons

We performed a set of behavioral experiments to investigate whether post-injury-born neurons also contribute to functional recovery. To determine if killifish can learn to avoid an aversive stimulus, we first tested if killifish naturally prefer a dark or light zone in a maze. Fish were allowed to freely explore a maze consisting of equally sized dark and light arms, separated by a central zone, for 15 min. Across three independent experiments, an innate preference for the dark arm was consistently observed (Fig. 6A, two-tailed paired t-test, p_i_= 0.0222 (t=2.758, df=9), n_i_=10; p_ii_=0.0095 (t=3.314, df=11), n_ii_=12; p_iii_= 0.0115 (t=3.030, df=11), n_iii_=12, same cohort as ii but one day later). Next, we paired entry into the dark arm with a mild electric shock to assess aversive learning (Fig. 6B). Each time a fish entered the dark arm, a mild shock was delivered. Recordings lasted 20 minutes and were repeated for five consecutive days. Three out of eight fish gradually reduced the time spent in the dark/punish arm over the course of five days (Fig. 6Bi). We termed these fish *learners*. In contrast, fish that failed to reduce their time in the dark arm across sessions were termed as *non-learners*, as they continued to spend a substantial portion of the 20-minute test in the punish arm. Upon comparing the time spent in the punish arm, we detected a significant difference from the third day onward, with leaners spending less time in the dark arm compared to non-learners (p_D3_= 0.0017, p_D4,D5_<0.0001, Two-Way ANOVA, Fig. 6Bii). Two-way ANOVA revealed a main effect of day of testing, F (4, 30) = 4.990, p = 0.0033, a main effect of learning, F (1, 30) = 55.92, p < 0.0001, and a day of testing × learning interaction, F (4, 30) = 4.801, p = 0.0041. Next, a double bilateral injury was inflicted in the Dm zone of the telencephalon of the learner fish (Fig. 6C). This region is predicted to be the homolog of the mammalian amygdala in zebrafish and goldfish, and thus to be involved in fear/associative learning and memory [14]. We hypothesized that this would also impair memory retention and the ability to relearn the task in killifish. When retested between 1 and 4 days post-injury, the previously learned avoidance behavior was lost, and the fish again spent significantly more time in the dark (punish) arm (Fig. 6Ci). When comparing the fish before and after the injury, we detected a significant increase in the amount of time spent in the punish arm from day three onward for the lesioned fish, comparable to the time spent in the dark arm for non-learners (Fig. 6Cii, p_D3_=0.0029, p_D4_= 0.0015, two-way ANOVA). Two-way ANOVA revealed a main effect of day of testing, F (3, 16) = 5.679, P=0.0076, a main effect of injury F (1, 16) = 27.76, p < 0.0001, and a day of day of testing × injury interaction, F (3, 16) = 5.649, p = 0.0078. In a set of learner fish in which the injury was deliberately placed lateral from the Dm zone, or did not target the whole anterior-posterior axis of the Dm zone, or was inflicted only unilaterally, learning and memory capacity was not impaired. One to four days post injury, these fish were perfectly able to avoid the dark arm (Supplementary Fig. 6). This is an extra indication that the full anterior to posterior Dm zone, and not the more lateral Dc/Dl zone or a subpart of the Dm zone, is essential for effective avoidance learning and memory. Together, these observations indicate that, also in killifish, the Dm zone of the telencephalon is responsible and essential for avoidance learning and memory and thus homologous to the mammalian amygdala.

**Figure 6:**
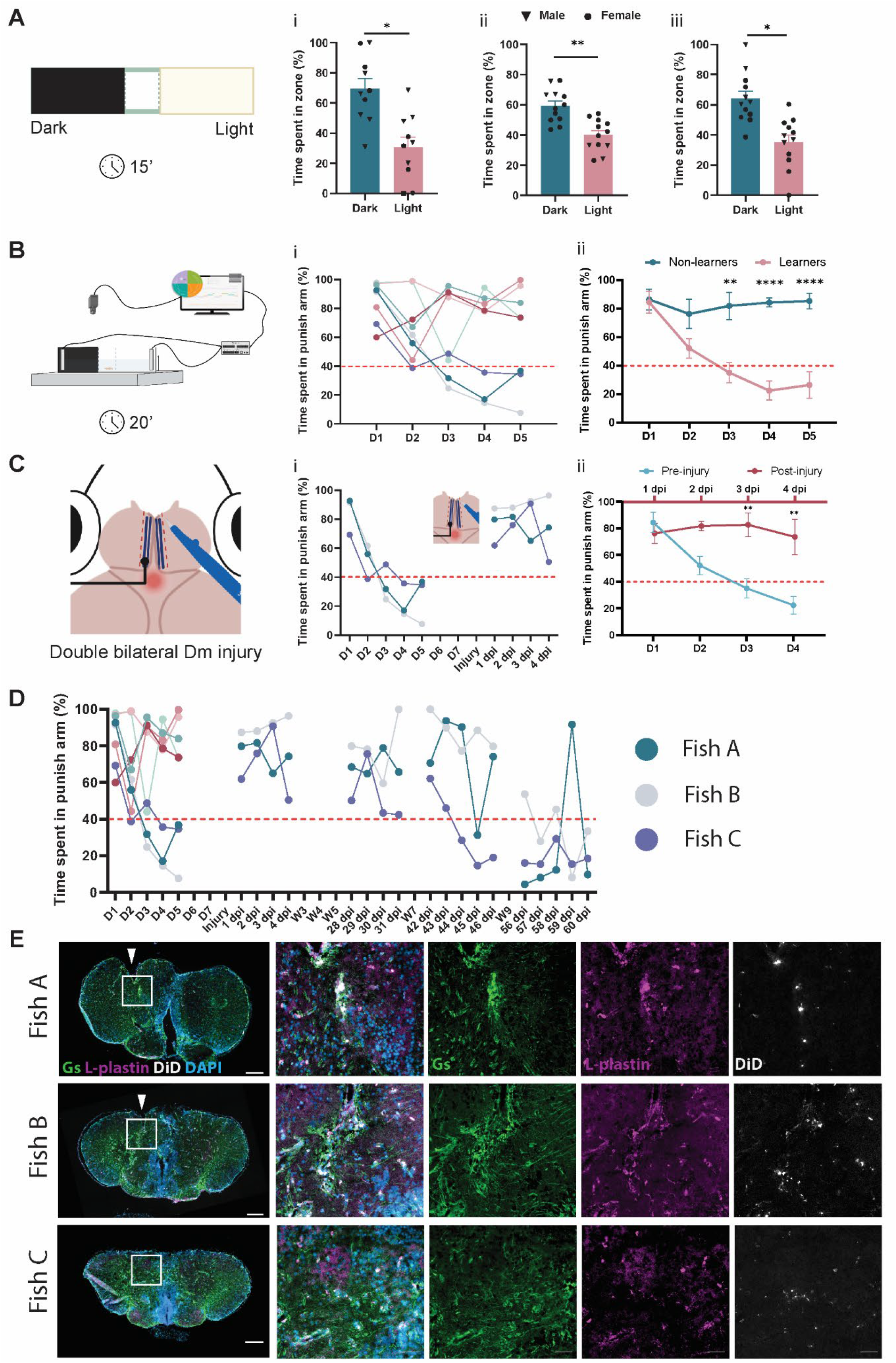
Behavioral readout of functional recovery. (A) Light-dark preference of the killifish is tested during 15 minute recordings. (i) The first cohort of 10 fish shows a preference for dark over light (center + light arm). (ii) A second cohort of 12 fish also shows a preference for the dark. (iii) A day later, the second cohort still shows the same preference for the dark, p_i_= 0.0222; p_ii_=0.0095; p_iii_= 0.0115. (B) Conditioned place avoidance experimental setup, where the dark arm of the maze is coupled to a mild electric shock. The fish are placed in the center of the maze and acclimated for 10 min. Next, the fish can freely explore the maze. Upon entering the dark arm, a shock is given for 3s. Whenever re-entering the dark arm, the same 3s shock is given. In the event the fish dwells in the dark arm, 3s-shocks are interleaved with 15s-pauses. (i) Over the course of 5 days, 3 out of 8 fish learned to avoid the dark arm. These fish are termed learners. (ii) When comparing learners and non-learners, already from the third day, a significant learning effect is observed. Mean with SEM, n_learners_ = 3, n_non-learners_ = 5. (C) A double bilateral injury is inflicted in the Dm zone of the telencephalon of learner fish. (i) The effect of the injury is tested by performing the conditioned place avoidance test. The fish show loss of memory and learning, and they no longer avoid the dark arm. (ii) Comparing pre- and post-injury learning abilities also shows that fish cannot relearn to avoid the dark arm as they could previously on day 3. Mean with SEM, n_pre-injury_ =3, n_post-injury_ =3. (D) Upon longer recovery from injury, fish were tested again in the conditioned place avoidance test. Recovery was expected by 30 dpi based on histology [12], but here the fish still spent most of their time in the dark arm. Fish were retested at 42 dpi. Fish C clearly relearned to avoid the dark arm in this week, while fish A and B still spent most of the time in the dark arm. By 56 dpi, fish C stayed in the light arm and fish B learned to do so in this week. (E) After behavioral testing, a staining for GS (Gluthamine synthetase – green), L-plastin (microglia – magenta), autofluorescent DiD (injury site – white) was performed. The brain of fish C shows no tissue malformation and minimal inflammation; while in fish A and B, clear signs of inflammation are visible as well as a malformation of the brain (arrowhead). Scale bar: 200 µm.

Purely based on histological analysis of wound healing and synaptic coverage, we expected recovery of behavior around 30 dpi [12]. Yet, at 30 dpi the learner fish still spent a large amount of the time in the dark arm (Fig. 6D). Retesting by 42 dpi revealed relearning in one out of three fish (Fish C – Fig. 6D). The other two fish still spent a lot of time in the dark area around 45 dpi. By 56 dpi all three fish displayed a decrease in the time spent in the punish arm and thus clear signs of functional behavioral recovery. We questioned if the observed difference in time post-injury needed for the aversive learning behavior to reappear was reflecting a different regenerative ability and thus a different speed of full brain tissue recovery. GS (Glutamine Synthetase, glial marker) and L-plastin (microglia marker) stainings indeed confirmed that fish C (fastest learner) had the lowest level of inflammation, since almost no L-plastin+ microglia/macrophages were visible, while the brains of fish A and B (slower learners) still showed more inflammation and residual tissue malformation (Fig. 6E - arrowheads). In addition, GS positive cell bodies and fibers can be detected in the malformed injury-site of fish A and B, as typically observed in the context of glial scarring [12], while less visible in the injury-site of fish C.

A second batch of fish was trained in an adapted version of the avoidance test in an attempt to increase the number of learners (Supplementary Fig. 7). On day 1 the fish were, as in experiment 1, placed in the center of the maze but now the fish with the strongest dark preference were selected (n = 8 out of 15, data not shown). Upon entering the dark arm, they received one electric shock for 3s to make the association between the dark arm and the electric shock. On day 2, these fish were placed in the same maze and allowed to swim freely, but now fish that dwelled in the dark arm for longer than 30s, despite the mild electroshocks, were placed back in the center zone and the trial restarted. Our prediction was that this might increase the fish’s correlation between the dark area and the electric shock. From day 3, the protocol was identical to experiment 1. Fish could freely swim for 20 min and they would receive a mild electric shock once entering the dark arm. Out of 8 fish, 4 learned to avoid the punish arm effectively with this approach. Upon receiving a double bilateral injury, 3 out of 4 fish recovered behaviorally around 45 dpi (Supplementary Fig. 7). The fourth learner could already perform the test from 1-4 dpi, where it gradually decreased the time spent in the dark arm over the 5 testing days (1dpi: 52%, 2dpi: 46%, 3dpi: 20%, 4dpi: 14%, 5dpi: 26%). In this fish, the injury turned out only to cover the anterior Dm zone (data not shown).

## 3 Discussion

Our study provides insights into how the short-lived killifish is an ideal model to study the mechanisms behind successful neuroregeneration. We found compelling evidence at multiple levels of inquiry that young adult killifish can recover functionally upon injury. The morphology of new neurons born after TBI is indistinguishable from naive neurons by 50 to 70 dpi, a bit slower than what is observed for constitutive growth and preceded by an overgrowth phase of maximal neurite branching. Relying on patch-clamp electrophysiology, we witnessed maturation of electrical properties, functional synaptic input and an abundance of dendritic spines from 50 dpi onward. Consistent with the timing of the maturation of all these morpho-electric cellular features, the regain of functional avoidance behavior in a conditioned place avoidance test occurred in a similar time frame. In sum, full recovery from brain injury in the Dm zone of the telencephalon occurs within approximately 50 days and involves the successful integration of adult-born mature neurons into the neural circuitry of the young adult killifish brain.

Mechanical brain injuries, such as the slit-injury implemented here, are widely used in regeneration studies in both mice and teleost fish [19–25]. This method remains a preferred approach since it does not require specialized equipment and the surgery can be done within 5 minutes. In our study, we specifically chose to injure the anatomically-defined Dm (Dorsomedial) zone of the killifish telencephalon [15] to enable behavioral assessment of functional recovery. Prior studies in zebrafish and goldfish indicated that the teleostean Dm region is homologous to the mammalian amygdala, based on its role in fear/avoidance learning and memory [26–28]. Following slit-injury, killifish failed to perform the learned task for several weeks. This response is similar to what is observed in goldfish after a Dm injury [26]. In contrast, lesions in a more lateral part of the goldfish pallium, the Dl zone, did not impair avoidance learning in a non-trace avoidance conditioning experiment [26]. Similarly, in our experiments, avoidance learning was preserved when the injury was placed lateral to the Dm, was unilateral, or did not fully extend along the anterior-posterior axis. These results confirm that the Dm region is critical for avoidance learning in killifish.

A notable limitation of the behavioral experiments was the low proportion of learners per trial, a phenomenon also reported in other teleost studies [26]. Variability in learning performance may be influenced by factors such as cognitive capacity, motivation, stress, and personality traits. For example, shy trouts have been shown to outperform bold individuals in maze-based food-reward tasks [29]. In future experiments, preselecting killifish based on personality traits may increase the proportion of learners. Recent work by Thoré and colleagues demonstrated that more active killifish exhibit lower risk-taking behavior and tend to stay near the periphery in open field tests [30]. Thus, selecting less active killifish may enrich for fish more inclined to take risks, such as escaping the dark arm after an electric shock, and thereby increasing the number of learners.

We observed variability in the speed and extent of regeneration among individual fish based on recovery of avoidance behavior, which might reflect interindividual differences in regenerative capacity arising from genetic or physiological variability. Variability in the original size of the injury might also play a role, with fish exhibiting larger lesions tending to show slower recovery. Notably, the fish with delayed functional behavioral regeneration also showed stronger signs of glial inflammation, which again may be a consequence of more extensive damage or a difference in individual aging rate since recovered fish approach the age of 16 weeks [12]. Histological analysis of the telencephalon after relearning delivered evidence that the speed and degree of relearning indeed corresponded with the physiological brain state.

Where in a vertebrate brain and how neurogenesis takes place appears to depend on two key factors: (1) the microenvironment, or neurogenic niche and (2) the cell intrinsic capacity for neurogenesis [31]. The killifish model allows studying both aspects, comparing the difference in microenvironment (injury vs. naive) as well as examining the cell-intrinsic properties of stem cells in these conditions or upon aging. When comparing adult born neurons in naive and injured fish, we observed a delay in maturation in the injured environment, potentially due to processes taking place in the microenvironment, such as local cell death, inflammation, and reactive progenitor cell proliferation. As shown in mice, the injury-induced microenvironment and inflammation can lead to aberrant synaptic functions, such as an increase in EPSC frequency and amplitude, and an increase in the number of spines [32,33]. Our previous study and our in situ sequencing assay (APOEB and PR2Y12) indeed demonstrated a high but acute inflammatory response early after injury in the telencephalon [12]. Overall, adult born neuron maturation and its timing appears to follow a cell-intrinsic program that is similar in both conditions. Still, we cannot exclude that in young killifish both microenvironments (naive and injured) are suitable to support successful neurogenesis. Hence, what would be interesting in the future is to transplant potent stem cells from young killifish into aged killifish brains and investigate the maturation of their progeny. The aged microenvironment is judged non-permissive due to a high burden of senescence and clear signs of chronic inflammation, and because cellular regeneration upon injury no longer occurs [12], yet the cell-intrinsic program of young proficient stem cells might be able to overcome these hurdles and still support effective neuron maturation even in an aged context. Vice versa, transplanted old stem cells might differentiate into mature neurons in the young killifish brain due to the permissive environment, or still fail to mature and integrate due to changes in the cell-intrinsic properties of aged stem cells. It has been shown before that the maturation and integration of neurons follow a cell-intrinsic program after transplantation of human neurons into the mouse cortex [34]. Human neurons only develop a full mature morphology over the course of one year in the mouse cortex, while for mouse neurons, this takes three to four months [32–34].

Retroviral labeling for lineage tracing of newborn neurons has been foremost used in mammals [35,37], and only sporadically in teleost fish [13]. In Killifish, our MLV strategy ensured the transduction of stem cells and the tracing of their progeny, creating sparse labelling of newly formed neurons, ideal for morpho-electric analysis. As expected, only a small portion of transduced cells were not of neuron identity and were located at or near the ventricular surface where the NGPs, radial glia and early-differentiating progenitors reside [12,17]. Based on our in situ sequencing data, we predict most of these non-neuronal cells to be NGPs since these are detected in larger numbers in the injured Dm zone at 2 dpi and they are in close proximity of cell division markers PCNA and MKI67. In addition, it is also known that the NGPs are responsible for the bulk of neuronal proliferation upon injury and during accelerated brain growth in the killifish telencephalon [10,12,18].

At the start of our investigation, we expected that regeneration after injury, and hence functional recovery, would take around 30 days, based on previous immunohistological observations about tissue healing and absence of glial scarring upon stab-wound injury in young adult killifish [12]. Our observation that the cell bodies and neurites of adult-born neurons were covered by presynaptic contacts at 23 dpi, suggestive of synaptic maturation, further triggered that expectation. However, behaviorally, fish still spent most of their time in the punish arm of the maze at 30 dpi. Only by 56-60 dpi, fish relearned to avoid the punish arm, which indicates that full functional recovery takes longer than overall structural recovery. At the cellular level, adult-born neurons still displayed too many neurite branch points around 30-36 dpi, indicating that an ensuing pruning phase is still necessary to reach full morphological maturity [38,39]. This pruning phase results in a net neurite structure that resembles that of naive mature neurons, just like adult-born neurons in mammals, and recapitulating developmental steps of neuron maturation [40]. This metabolically costly process may facilitate the selection of those synaptic partners that, in the end, efficiently steer the relevant behavior.

Our observations thus are very reminiscent to those features witnessed in neurogenesis in mammals. Whole-cell recordings of birth-dated GFP+ neurons revealed larger spontaneous EPSC amplitudes in cells younger than 50 dpi compared to older or naive neurons, suggesting that the excitatory– inhibitory balance continues to be refined beyond this point. Similar to mammalian brain development, where immature neurons display high EPSC amplitude variability and spontaneous network activity guides synaptic pruning and stabilization [38,39,41,42], injury-induced adult-born neurons in killifish show heightened excitatory input early on, followed by maturation-related amplitude normalization by 50 dpi. This process is accompanied by a lower input resistance and increasing dendritic spine density, paralleling observations in human cortical and adult-born mouse hippocampal neurons [5,34]. This again shows the strength of the killifish to model mammalian processes regarding neurogenesis, which is very promising to find therapies to boost neuroregeneration.

In summary, we demonstrate that young killifish are capable of functional recovery following a slit-injury in the telencephalon, as evidenced by molecular, cellular and behavioral measures. Neuroregeneration takes around 50 days, which is relatively long for a fish with a short lifespan of only 4 to 6 months, raising questions on the scalability of regeneration between species. With this timeline for functional regeneration, it closely resembles the mammalian context, as in mice functional regeneration also takes up to 8 weeks [34–36]. The maturation process of neurons thus appears uncoupled from lifespan. This study also advances the field of killifish brain neuroregeneration research by expanding the toolbox with novel techniques, including viral vector labeling via ventricular and in-brain injections, electrophysiological recordings, and the establishment of a new behavioral assay for learning and memory. The field can now embrace this model and the optimized techniques and expand screening strategies for therapeutic compounds for improved neuroregeneration and neuronal maturation and integration in disease or aged conditions.

## 4 Materials and methods

### 4.1 Animals and housing

All experiments involved adult (6-week) male (light-dark behavior only) and female African turquoise killifish (*Nothobranchius furzeri*) from the GRZ-AD strain. This age was defined before based on a lifespan curve in our facility [12]. Fish were bred in 8L aquaria enriched with a sandbox for spawning. Experimental fish were housed in 3,5 L aquaria in a ZebTec multi-linking system. Three females were housed with one male, and they were fed twice a day with *Artemia salina* and Chironomidae mosquito larvae (Ocean Nutrition). Housing conditions were standardized: a 12 h/12 h light-dark cycle, a water temperature of 28°C, a water conductivity of 600 µS, and a pH of 7. For viral vector injections, three fish were housed in a 1L box for five days in an L2 environment in an incubator with daily water changes. For the first three days, the temperature was set to 36°C to enhance viral vector expression [43]. All experiments were approved by the KU Leuven ethical committee in accordance with the European Communities Council Directive of September 22, 2010 (2010/63/EU) and the Belgian legislation (KB of May 29, 2013).

### 4.2 Slit-injury

To study the regeneration process, fish were injured in the Dm zone of the telencephalon, as defined anatomically [15]. First, fish were anesthetized in 0.03% buffered tricaine (MS-222, Sigma Aldrich) diluted in aquarium system water and placed in an ice-cold, moist sponge under a stereomicroscope. Using fine forceps (FST, Dumont #5 11251-20), the scales and skin on top of the brain were removed as described before [12]. A micro knife was used to perform the injury from rostral to caudal (Fine Science Tools, 15° cutting angle, 10315-12). To anatomically standardize the injury depth, only 1 mm of the knife blade was used to injure the right telencephalic hemisphere, and the success rate for hitting the right brain area was around 70% (Supplementary Fig. 1). Fish used for histological analyses, received a single unilateral injury into the Dm zone of the right hemisphere of the telencephalon. For behavioral experiments, a double bilateral injury was inflicted into the Dm regions of the telencephalon. Fish recovered in fresh system water before returning to their home tanks. Before injuring the brain, the knife was dipped into Vybrant DiD cell-labeling solution (V22887, Thermo Fisher Scientific), leaving a visible trace of the injury site, even upon complete recovery. DiD crystals stick to the cell membranes upon injury and are auto-fluorescent (Ex: 644 nm, Em: 665 nm). In electrophysiology experiments the DiD crystals were not used to avoid bleed through during recording of GFP+ cells. The standardization of the injury was determined by measuring the depth of the injury and the distance to the midline using Fiji.

### 4.3 EdU labeling experiment

After anesthesia in 0.03% tricaine, 10 µL of 5 mM 5-ethynyl-2′-deoxyuridine (EdU, Invitrogen, C10340) was injected intraperitoneally (i.p.) using a 33-Gauge Hamilton Syringe. EdU injections were administered every morning at 8 a.m. and every evening at 6 p.m. for 4 consecutive days to allow constant labeling of dividing cells. At 12 p.m on day 1, the retroviral vector was injected into the ventricle, and half of the cohort of fish also received an injury in the Dm zone (n=3). After a chase of 21 days, brains were collected, sectioned and stained for GFP and EdU (Fig. 1A, B, Supplementary Fig. 2).

### 4.4 Viral vectors

The retroviral vector (ɣ-retroviral, murine leukemia virus, MLV-based, EFs-eGFP, TU/mL = 1,30E+08 determined on HEK293T cells) used to trace the progeny of dividing progenitors was produced using tiple transient transfection by the Leuven Viral Vector Core (LVVC; www.lvvc.be) and expresses eGFP under the EFs (human elongation factor 1 α short) promoter as previously described [44]. The recombinant adeno-associated viral vector (AAV) exclusively used for Sholl analysis of neurons in naive fish expresses eGFP under control of the CAG promoter (Addgene, 37825-AAV1, titer ≥ 7×10^12^ vg/mL). Viral vectors were always injected at a maximal titer, with the addition of 10% fast green (Sigma, F7252-5G) to visualize the injection spread into the ventricle. Successful injection could be confirmed by the spread of the green color between the two telencephalic hemispheres [45]. The viral vector was either injected in the brain parenchyma directly or in the telencephalic ventricle, as explained below.

#### 4.4.1 Cerebroventricular injections

Cerebroventricular injections were carried out according to the injection protocol for zebrafish of Ninkovic and Durovic [45] to label progenitor cells and their progeny. In short, fish were anesthetized in 0.03% tricaine and placed in a cold, moist sponge with the dorsal side facing up. The scales on top of the head were removed and the skin was opened with two forceps (FST, Dumont #5 11251-20), after which the skull became visible. Using a micro knife (FST, 10315-12), a small slit was made in between the telencephalon and the optic tectum as an opening for the injection capillary. Using a FemtoJet 4i (Eppendorf) and a micromanipulator, the capillary was placed through the opening in between the two telencephalic hemispheres, and a volume of max. 0.5 µl was injected. The fish were placed in fresh system water to recover and were housed in an L2 environment for five days. The first three days, the fish were housed at 36°C, from then on again at 28°C, since this has been reported to increase viral vector expression in zebrafish [43].

#### 4.4.2 In-brain injections

Injections of an AAV2.1 vector into the brain parenchyma were performed to label mature neurons in the Dm zone of naive killifish for Sholl analysis. A Nanoject II microinjector (Drummond Scientific) was used to deliver ±100 nL of the AAV vector into the brain tissue. Fish were anesthetized in 0.03% tricaine and placed in a cold, moist sponge with the dorsal side up. The scales were removed on top of the telencephalon and the skin was opened until the telencephalon was visible using two forceps (FST, Dumont #5 11251-20). A small piece of the skull was removed with fine forceps (FST, Dumont #5 11251-20) and the capillary containing the vector was inserted using a stereotactic frame to a depth of approximately 300 µm. 92 nL was injected in steps of 4 x 23 nL and after injection the capillary remained in the brain tissue for another minute to allow the spread of the viral vector. The fish were placed in fresh system water to recover and were housed in an L2 environment for five days. The first three days, the fish were housed at 36°C and then returned to 28°C.

### 4.5 Tissue fixation and processing

At a specific time point after injury (11, 18, 21, 23, 24, 30, 36, 50, and 70 days post-injury), fish were perfused intracardially to preserve tissue morphology as described previously [46,47]. Fish were killed in 0.1% buffered tricaine diluted in aquarium system water, followed by perfusion through the heart with 1X PBS followed by 4% paraformaldehyde (PFA, in PBS, Sigma-Aldrich). The brain was collected and post-fixed overnight in 4% PFA. Next, the brain was washed three times with 1X PBS and embedded in either 4% agarose (vibratome sections) or saturated with 30% sucrose and embedded in 1.25% agarose, 30% sucrose in PBS (cryosections). In case of cryosectioning, 10 µm-thick sections were made on a cryostat (Leica CM3050s), collected on Superfrost Plus Adhesion Slides (Thermo Fisher Scientific), and stored at -20°C. For vibratome sections, 50 µm-thick sections were made on a vibratome (HM 650 V, Microm Microtech), collected and stored in 24-well plates at 4°C until further use.

### 4.6 Whole brain explant preparation

Fish were killed with an overdose of oxygenated tricaine (0.1%), decapitated and the head placed in ice-cold oxygenated (95% O_2_/5% CO_2_) artificial cerebrospinal fluid (ACSF; 124 mM NaCl, 22 mM D-glucose, 2 mM KCl, 1.6 mM MgSO_4_.7H_2_O, 1.25 mM KH_2_HPO_4_, 24 mM NaHCO_3_, 2 mM CaCl_2_.2H_2_O; 310 mOsm, pH 7.4). The lower jaw, the eyes and all the remaining bone and tissue surrounding the brain were then removed. Once the brain was isolated, the optic nerves were cut using forceps (FST, Dumont #5 11251-20), and the *tela choroidea* was removed with very fine forceps (Dumont #5 – Fine forceps – 11254).

### 4.7 Electrophysiology

Electrophysiological recordings were carried out on *ex vivo* whole brain explants, which were placed in a recording chamber and kept in place using a fine-wired slice anchor (Warner Instruments, SHD-26GH/10). A continuous flow of RT oxygenated ACSF (2ml/min) was maintained by using a peristaltic pump (PPS2, Multichannel systems). Borosilicate glass micropipettes were used (Science Products, GB150F-10P) and pulled (P-97, Sutter Instrument) to create resistances between 6-9 MΩ. The intracellular solution contained 10 mM KCl, 10 mM HEPES, 4 mM NaCl, 4mM Mg-ATP, 10 mM phosphocreatine, 0.3 mM Na-GTP, 130 mM K-gluconate with an osmolarity of 295 and pH of 7.2. Biocytin was added in a concentration of 0.2% to visualize the morphology of the recorded cells. Newborn neurons were visualized using an upright microscope (Axio Examiner.A1, Carl Zeiss NV, Belgium) equipped with a 5X, 0.16 NA and a 40X 1.0 NA objective and complemented with a fluorescence LED illumination system (Colibrí 5, Zeiss) to identify their GFP fluorescence. Recordings were amplified and digitized using an integrated patch amplifier (IPA, Sutter Instruments), and analyzed off-line either with Clampfit (Molecular Device, v.11.2) (IV protocol, gapfree VC) or IgorPro (SutterPatch sofware) (CC step, gapfree CC). Cells were rejected if the input resistance was below 0.3 GΩ, if the resting membrane potential was more depolarized than -30 mV, and if the series resistance was higher than 20 MΩ. Maximal time to patch cells after dissection of the brain was three hours. Cells were recorded for maximally 30 min to maintain the cell body in the brain for biocytin reconstruction. To this end, the pipette was slowly retracted from the cell by moving away diagonally.

### 4.8 Immunohistochemistry

Vibratome sections were washed in PBS and mounted on gelatin-coated glass slides. After drying for 30 min at 37°C for adhesion, the slides were washed 3 times for 5 min in PBST (0.1% Triton-X-100). Blocking was performed by incubating the slides for 2 h at RT in 20% pre-immune goat serum (Sigma-Aldrich, S26) in Tris-NaCl blocking buffer (TNB). The primary antibody was applied overnight in TNB at RT or in Pierce Diluent for the HuCD antibody. Primary antibodies used were chicken anti-GFP (1:2000, Abcam, ab13970) mouse anti-HuCD (1:100, Thermo Fisher Scientific, A-21271), rabbit anti-lcp1 (1:1000; Sigma, SAB2701743), mouse anti-Gs (1:1000, Abcam, ab64613). After overnight incubation, the slides were washed three times with PBST (0.1% Triton-X-100). Secondary antibodies were applied at RT in TNB for 2 h. For anti-L-plastin (Lcp1) IHC staining, an amplification was used, in which the secondary antibody is coupled to biotin (Goat anti-Rabbit-biotinylated, 1:300 in TNB, E043201-8, Agilent Dako) and applied for 45 min. After washing three times for 5 min with PBST (0,1% Triton-X-100), Streptavidin-594 (1:500 in TNB, Thermo Fisher Scientific, S32356) was added to the sections for 2 h. Other secondary antibodies used were Goat anti-chicken Dylight 488 (1:300 in TNB, Abcam, ab96947) and Goat anti-mouse 488/594 (1:300, Invitrogen, A11001/A-11005). Nuclear staining was performed using 4’,6-diamidino-2-fenylindool (DAPI, 1:1000 in PBS, Thermo Fisher Scientific). Slides were mounted with Mowiol/SlowFade Glass (Life Technologies, S36920) and coverslipped.

Full brains from electrophysiology were fixed overnight and thereafter washed with PBST (0.5% Triton-X-100), three times for two hours. Blocking was performed by incubating for 2 h at RT in 20% pre-immune goat serum (Sigma-Aldrich, S26) in PBST (0.1% Triton-X-100). The primary antibody (chicken anti-GFP (1:2000, Abcam, ab13790) was applied overnight in PBST (0.1% Triton-X-100) at RT. After overnight incubation, the brains were washed three times with PBS (0.1% Triton-X-100). Secondary antibodies were applied overnight at RT and were Streptavidin-Cy5 (1:500 in PBST (0.1% Triton), Invitrogen, SA1011) and Goat anti-chicken Dylight 488 (1:300 in PBST (0.1% Triton), Abcam, ab96947). Nuclear staining was performed using DAPI. The brains were mounted in imaging solution (4% gelatin and 65% glycerol in PBS) and coverslipped.

### 4.9 Microscopy

Images were acquired with a widefield (Axio Observer7, Zeiss) or confocal (LSM 900 with Airyscan 2, Zeiss) microscope using Plan-Apochromat objectives 20X/0.8, 40X/0.95 or 63X/1.4. The images were further processed with ZEN 3.7 software (Zeiss). Z-stack images of cells after biocytin filling in the whole brain were acquired with a 20X objective in steps of 0.44 microns per plane, 2048 x 2048 px, 8 bit, zoom 0.7, and averaging 4. Images of spines were made using Airyscan mode, a 63X objective in steps of 0.19 micron, 1013 x 1013 px, 16 bit, zoom 2.0, and averaging 4. Image 1E has been created using image registration. The GFP images was taken using a Zeiss Axio Observer, and thereafter the slice was stained for HuCD and a second image was made. Image registration was performed using Fiji with descriptor based registration.

### 4.10 Sholl analysis

Z-stacks were acquired on the confocal microscope using the 20X/0.8 objective. Individual plane images were exported as TIF files and imported in FIJI [48]. The images were used to generate a stack and these were traced using the Neuroanatomy Plugin [49]. Following tracing, Sholl analysis was carried out with the cell body as center node using the SNT plugin in FiJi [49].

### 4.11 Conditioned place avoidance

A custom plexiglass maze (rectangle, 50x10x10 cm) was used, consisting out of a central habituation zone (10x10 cm), a dark arm (20x10 cm) and a light arm (20x10 cm) (Fig. 6). The dark zone was made by placing a black infrared-translucent plexiglass sleeve around that zone of the maze. The maze was placed on an infrared backlight, and recordings were performed with a GigE camera and the Ethovision XT 17 software (Noldus). The maze was filled up to 5 cm with aquarium system water, which was changed between every recording as described previously [50]. In the light-dark preference test, fish were allowed to swim freely for 20 min. For the conditioned place avoidance tests, two copper electrodes were added to the setup, one on the left and one on the right side of the maze (light and dark arm) and connected to an aversive stimulator (ENV410-C, Med Associates Inc.) that was controlled via the trial hardware control of the Ethovision Software. A mild electric shock of 3.5 mA was given, at which a consistent behavioral response was observed. In conditioned place avoidance test 1, fish were acclimated in the center of the maze for 10 min, after which the gates to the light and dark arms were opened. If the fish entered the dark arm, an electric shock was given for 3 s. If the fish did not escape the dark arm, the shock was repeated every 15 s. The total trial time was 20 min. The fish were tested individually for five consecutive days between 10 a.m. and 4 p.m., and the time spent in the dark arm was measured. Fish were termed learners if they spent less than 40% of the time in the dark arm on the last two testing days. The fish that spent most of their time (>40%) in the dark arm were termed non-learners. Learners were given a double bilateral injury in the Dm zone and were tested again from 1 to 4 days post injury (dpi) to interrogate if memory was lost due to the Dm injury. To evaluate re-learning and thus regeneration, the fish were tested again in three 5-day testing periods, at 28-32 dpi, 42-46 dpi, and 56-60 dpi. Hereafter, fish were perfused and brains collected for further histological analysis. In a second independent learning paradigm, small adaptations to the protocol were implemented to try to increase the number of learners in a batch of fish. Fish were tested on day 1 for their dark preference and received one electric shock for 3s to learn to correlate the dark arm with the electric shock. Fish with the shortest latency to reach the dark arm were selected for testing in the maze on day 2. On day 2, in case a fish remained in the dark zone for longer than 30 seconds, the trial was stopped and the fish was placed back into the center zone. The trial was then restarted. A fish was maximally placed back in the center 15 times. From day 3, fish were allowed to freely behave in the maze as in the first learning paradigm. In learning paradigm 1, 3 out of 8 fish were able to learn to avoid the dark arm, while in learning paradigm 2, 4 out of 8 fish were able to learn.

### 4.12 In situ sequencing

Telencephali of 6w naive and 6w-2dpi fish were collected after the fish were killed using an overdose of Tricaine (0.1%). The telencephalon was snap-frozen in cold (-50 °C) isobutane and stored at -80°C until sectioning. The telencephalon of 6w naive (n=3) and 6w-2dpi (n=3) fish was cryo-sectioned at 10 µm thickness in Tissue-Tek O.C.T. Compound (Sakura) and one anterior, one middle and one posterior section was collected on a Superfrost Plus^TM^ Adhesion Slide (Thermo Fisher Scientific; 10149870). In situ sequencing (ISS) was performed at the In Situ Sequencing Facility of SciLifeLab, Sweden, following established protocols [51,52]. For this study, padlock probes were designed to target a panel of indicated genes (Supplementary Table 1). A detailed step-by-step protocol for the ISS chemistry can be found in [53]. Briefly, 10 µm-thick cryosections were fixed in 3% (w/v) paraformaldehyde (PFA) in PBS at RT for 10 min, washed with PBS, and treated with 0.1 N HCl for 5 min at RT. After a subsequent PBS wash, the sections underwent reverse transcription, padlock probe ligation, and rolling circle amplification (RCA). RCA products were detected by hybridization and decoded through cyclic imaging rounds to identify the targeted genes. Each cycle involved base-specific incorporation of fluorescent dyes (A = Cy5, C = AF750, G = Cy3, T = AF488), followed by imaging and removal of the incorporated dyes. Fluorescence images were acquired using a Zeiss Axio Imager Z2 microscope (Zeiss, Germany) with a 40× objective. Illumination was provided by external LED sources (Lumencor SPECTRA X), and emission filters included filter paddles (395/25, 438/29, 470/24, 555/28, 635/22, 730/40). Images were captured on sCMOS cameras (2048 × 2048, 16-bit, ORCA-Flash4.0 LT Plus, Hamamatsu). Wavelength separation was achieved using filter cubes: quad band Chroma 89402 (DAPI, Cy3, Cy5), quad band Chroma 89403 (AlexaFluor750), and single band Zeiss 38HE (AlexaFluor488). For each sample, a series of images with 10% overlap between neighboring tiles was acquired, consisting of 28 z-stack planes at 0.3 µm intervals. Maximum-intensity projection (MIP) was applied to merge the stacks using Zeiss ZEN software. Exported images in.tif format were preprocessed for alignment between cycles and stitching of tiles. The preprocessing code is available at https://github.com/Moldia/ISS_preprocessing. Transcript decoding was performed using the Python package starfish (https://spacetxstarfish.readthedocs.io/en/latest/), and the decoding scripts can be accessed at https://github.com/Moldia/ISS_decoding. The decoded data were exported as a CSV file, providing an x–y coordinate map for each signal in combination with DAPI staining. The image data were visualized using TissUUmaps (https://tissuumaps.github.io/). We analyzed the following genes: HMGB2a (NGP marker), GAP43 (neurite outgrowth marker), MKI67 (proliferation marker), PCNA (proliferation marker) for their importance in neurogenesis, NEUROD2 to detect early differentiating neurons (Intercell-NC), APOEB, and P2RY12 for MG/MF, VIM- and GLUL for radial glia (RG1,2)

### 4.13 Statistical analysis

Statistical analysis was performed using GraphPad Prism v9.3.1. Results are shown as mean ± standard error of the mean (SEM). Data was first tested for Gaussian distribution. If assumptions were met, a two-tailed unpaired t-test (2 conditions) or one-way ANOVA (>2 conditions) followed by Dunnett’s multiple comparisons test was used. If assumptions were not met, a non-parametric two-tailed Mann Whitney test (2 conditions) or Kruskal-Wallis test (>2 conditions) followed by Dunn’s multiple comparisons test was used. A two-way ANOVA was used to compare the naive vs injured fish at the different timepoints post-injury. The number of data points and the exact test can be found in the figure legends. A cut-off of p ≤0.05 was used as the threshold for statistical significance.

## Author contributions

V.M.: Conceptualization, design, experiments, statistical analysis, writing, original draft, statistical visualization. C.Z.: review and editing. J.V.H.: Conceptualization, review and editing. A.M.: experiments, sholl analysis, review and editing. C.Z.: review and editing. R.A.: review and editing. C.V.D.H.: viral vector production, review and editing. R.G.: viral vector production, review and editing. M.T.: Help with electrophysiology experiments, review and editing. L.A.: Conceptualization, design, writing, review, editing, study supervision.

## Acknowledgements

We acknowledge the in situ sequencing facility, part of the Spatial Biology Platform at Science for Life Laboratory/Stockholm University, Sweden, for technical assistance with the in situ sequencing experiments.

