## Supplementary Figures for "From injury to recovery: functional neuronal regeneration after traumatic brain injury in the telencephalon of the young adult killifish"

Valerie Mariën[1], Caroline Zandecki[1], Jolien Van houcke[1], Anouk Maes[1], Rajagopal Ayana[1], Chris Van den Haute[2,3], Rik Gijsbers[2,4,5], Marialuisa Tognolina[1], Lutgarde Arckens[1,6*]

[1] Laboratory of Neuroplasticity and Neuroproteomics, Department of Biology, KU Leuven, Leuven, Belgium
[2] Leuven Viral Vector Core, KU Leuven, 3000, Leuven, Belgium
[3] Research Group for Neurobiology and Gene Therapy, Department of Neuroscience, KU Leuven, Leuven, Belgium
[4] Advanced Disease Modelling, Targeted Drug Discovery and Gene Therapy (ADVANTAGE), Department of Pharmacological and Pharmaceutical Sciences, Faculty of Medicine, KU Leuven, 3000 Leuven, Belgium
[5] Leuven Institute for Rare Diseases, Leuven, Belgium
[6] Leuven Brain Institute, Leuven, Belgium

**Corresponding author*

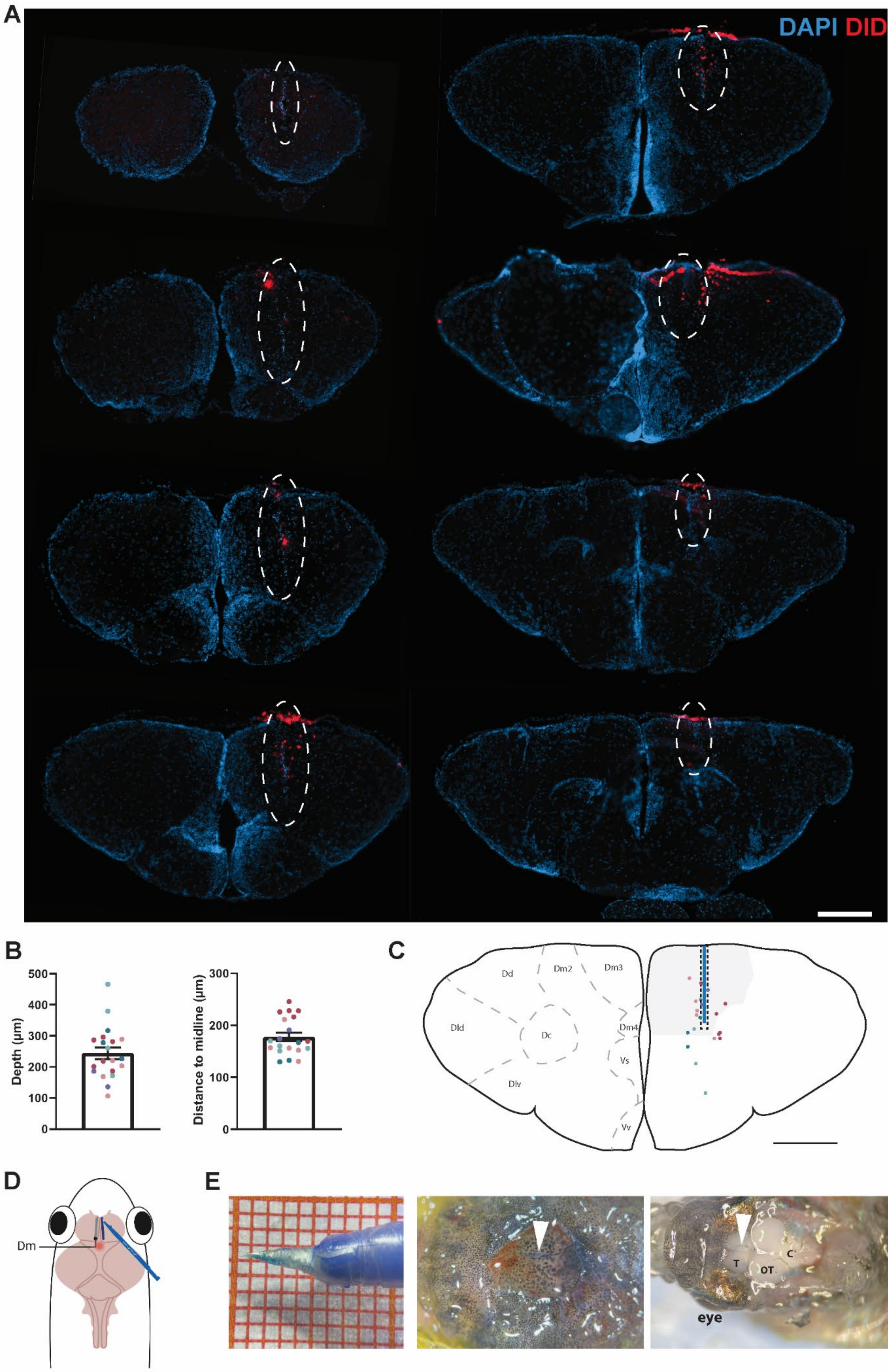

A
DAPI DID
B
Depth (µm)
500
400
300
200
100
0
Distance to midline (µm)
300
200
100
0
C
Dd
Dm2
Dm3
Dld
Dc
Dm4
Vs
Dlv
Vv
D
Dm
E
T
OT
C
eye

**Supplementary Figure 1: Standardization of the slit-injury in the anatomically defined Dm zone of the young adult killifish telencephalon.** (A) DAPI (blue) staining and auto-fluorescent signal of the Vybrant DiD cell labeling solution (red) on 10 µm-thick coronal telencephalic cryosections of a young adult (6 weeks) killifish at 1 day post injury (dpi). The injury is visible from rostral (upper left) to caudal (lower right) as indicated by dotted lines. Scale bar: 200 µm. (B) Quantification of depth of the injury and distance from the midline. Quantification was performed using ImageJ (FiJi). Error bars indicate the standard error of the mean (SEM). Color-coded dots correspond to different sections of one fish. Five fish were quantified and only when the injury was clearly measurable. (C) Scheme of a coronal section of the telencephalon, on the left hemisphere the different anatomical subregions are indicated based on the killifish brain atlas (D'angelo, 2013). On the right hemisphere the Dm zone is indicated in grey (Dm2-4), color-coded dots correspond to the different measurements as analyzed in B. The blue line represents the mean distance from the midline and mean depth, and the grey dotted line indicates the standard error of the mean. Scale bar: 200 µm. (D) Top-view schematic of the location of the Dm zone in the telencephalon, where the slit-injury is inflicted. (E) Left: the micro-knife is dipped in DiD dye and the blade depth used (part without parafilm) is approximately 1 mm. Middle: Blue-colored line from the DiD dye on the skull as the outcome of the injury. Right: after perfusion, this blue line remains visible in the Dm zone of the telencephalon. T: Telencephalon, OT: optic tectum, C: cerebellum.

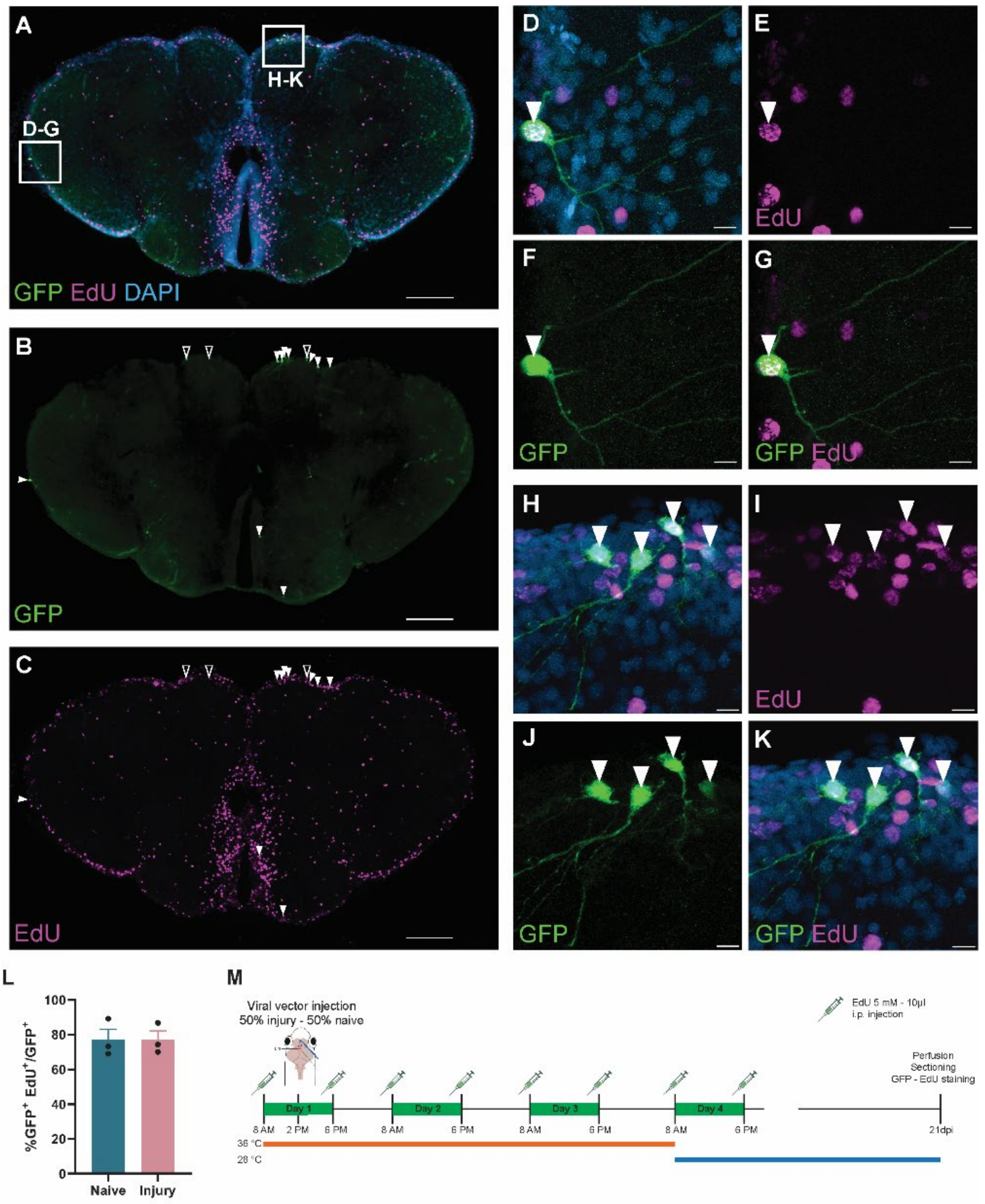


**Supplementary Figure 2: The ɣ-retroviral MLV vector specifically integrates into dividing cells.** (A-C) Coronal section of a naive killifish telencephalon stained for GFP (green), EdU (magenta) and DAPI (blue) 21 days post ventricular injection. 20X Tile image with white arrowheads pointing to double positive (GFP+/EdU+) cells and open arrowheads pointing to single GFP+ cells. (D-G) 63X Orthogonal projections of the boxed area indicated in A, showing a double positive newborn cell with long processes. (H-K) 63X Orthogonal projection of the boxed area indicated in A. (I) A gradient of EdU signal can be observed in the different GFP+ cells. (L) Quantification of GFP+/EdU+ cells in naive and injured fish at 21 dpi. All coronal sections (n=13-20) of the telencephalon were counted for three different fish per condition and a mean double labeling of 77% was observed in both conditions. (M) Scheme of the experimental setup. EdU was injected twice daily for four consecutive days. On the first day, fish were injected with the viral vector and half of the fish received an injury. The first three days, fish were housed at 36°C and from day four the temperature was lowered to the standard housing temperature of 28°C (see materials and methods). At 21 dpi, fish were perfused, brains were sectioned and the staining for GFP and EdU was carried out. Scale bar: A-C: 200 µm; D-K: 10 µm.

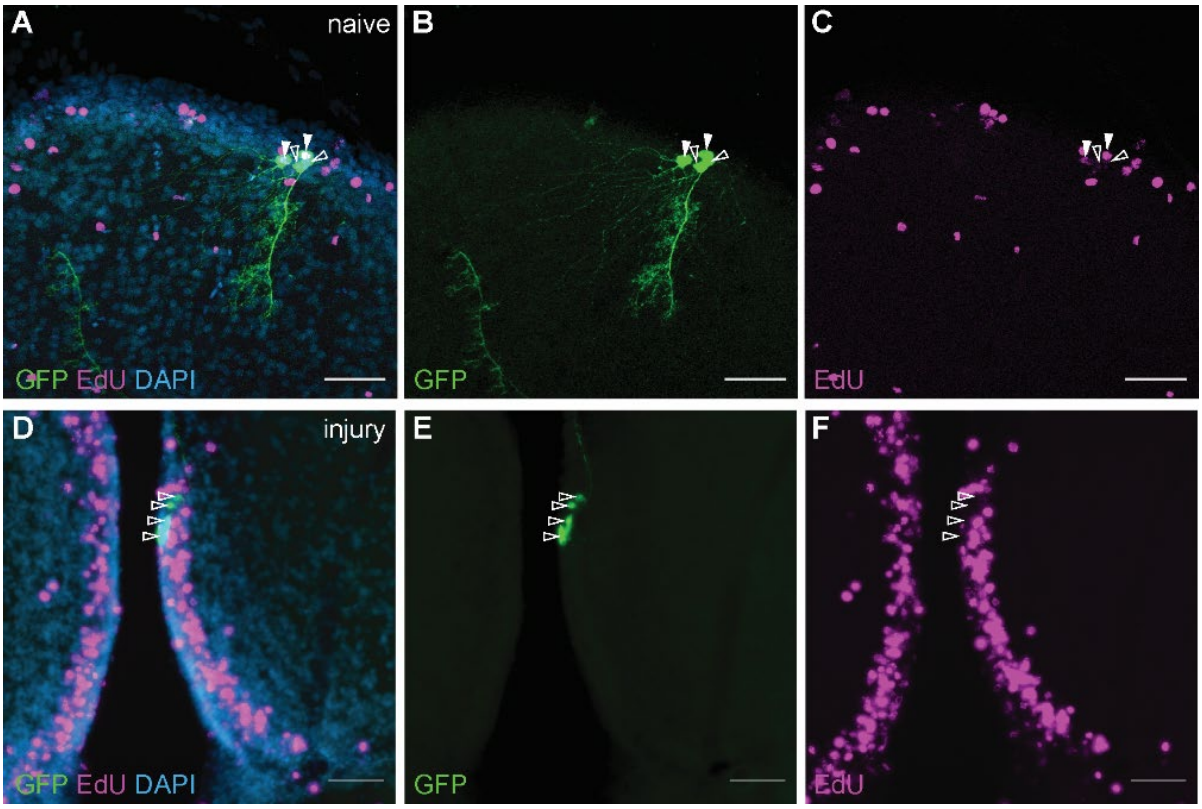


**Supplementary Figure 3: ɣ-retroviral MLV transduced cells that are EdU negative, are always located close to the ventricular zone, stem cell niche I and II** [10]**, and have a morphology similar to EdU+ transduced cells.** It is comprehensive that these ventricular cells lost EdU due to too many consecutive cell divisions. A-C: Example of GFP+ newborn cells in a naive fish, with some cells showing loss of EdU signal (open arrowheads). (D-F) Example from an injured fish at 21 dpi showing loss of EdU in all visible GFP+ cells. Confocal images taken with a 20X objective. Scale bar: 50 µm.

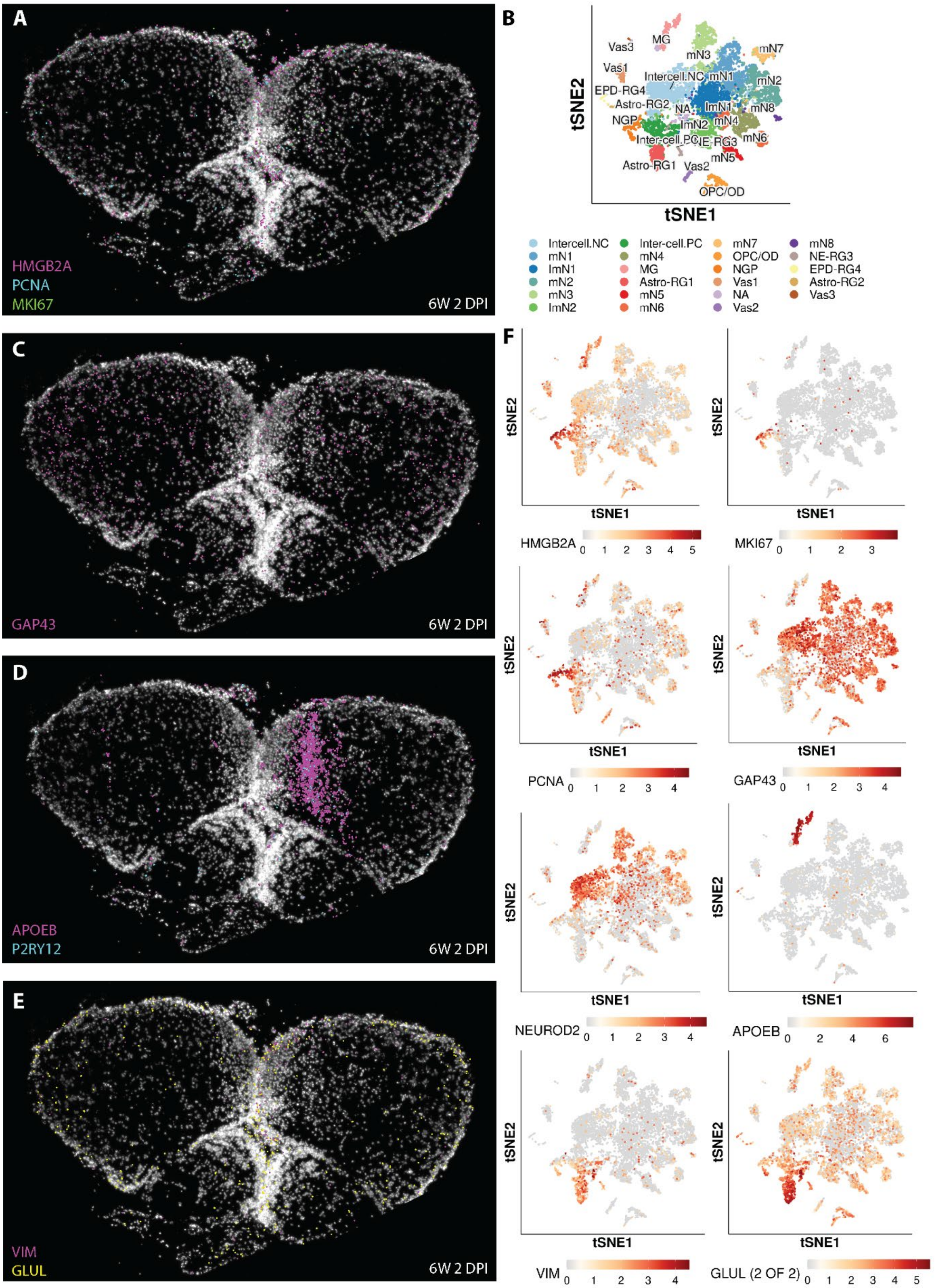


**Supplementary Figure 4: Shortly after injury (2dpi), in situ sequencing analysis reveals an increase in dividing NGPs (MKI67, PCNA and HMGB2A), along with increased expression of the axonal growth marker (GAP43), RG markers (VIM, GLUL) and a strong inflammatory response (APOEB and P2RY12).** (A, C, D, E) These increased signals characterize the injured Dm zone of the telencephalon (right hemisphere) more strongly than the contralateral hemisphere at 2 dpi. (A) Markers for cell division and NGPs are shown, overlap and are increased in the injured hemisphere. (B) TSNE plots of our previous single cell sequencing analysis shows 23 different cell populations, including those relevant to this study: MG, RG, IC-NC, NGP [17] (https://ayana-rajagopal.shinyapps.io/shinyappmulti_alls1/). (C) GAP43 expression is increased compared to naive fish and is more pronounced in the injured hemisphere. (D) Microglia/macrophages infiltrate the injury site at 2 dpi, as shown by the markers APOEB and P2RY12. (E) RG markers VIM and GLUL are increased in the injured Dm zone compared to the contralateral Dm zone. (F) TSNE plots for the different marker genes in panels A, C-E underscore the cell-type specificity of the probes used in the spatial seq analysis. HMGB2A, PCNA and MKI67 are typical markers for NGPs, GAP43 is found in early (Intercell-NC) and late (ImN) maturing and mature neurons (mN), NEUROD2 is mainly found in those intercells devoted to become neurons (intercell.NC), APOEB is specific to microglia, VIM and GLUL (Gs) are highly expressed by RG1.

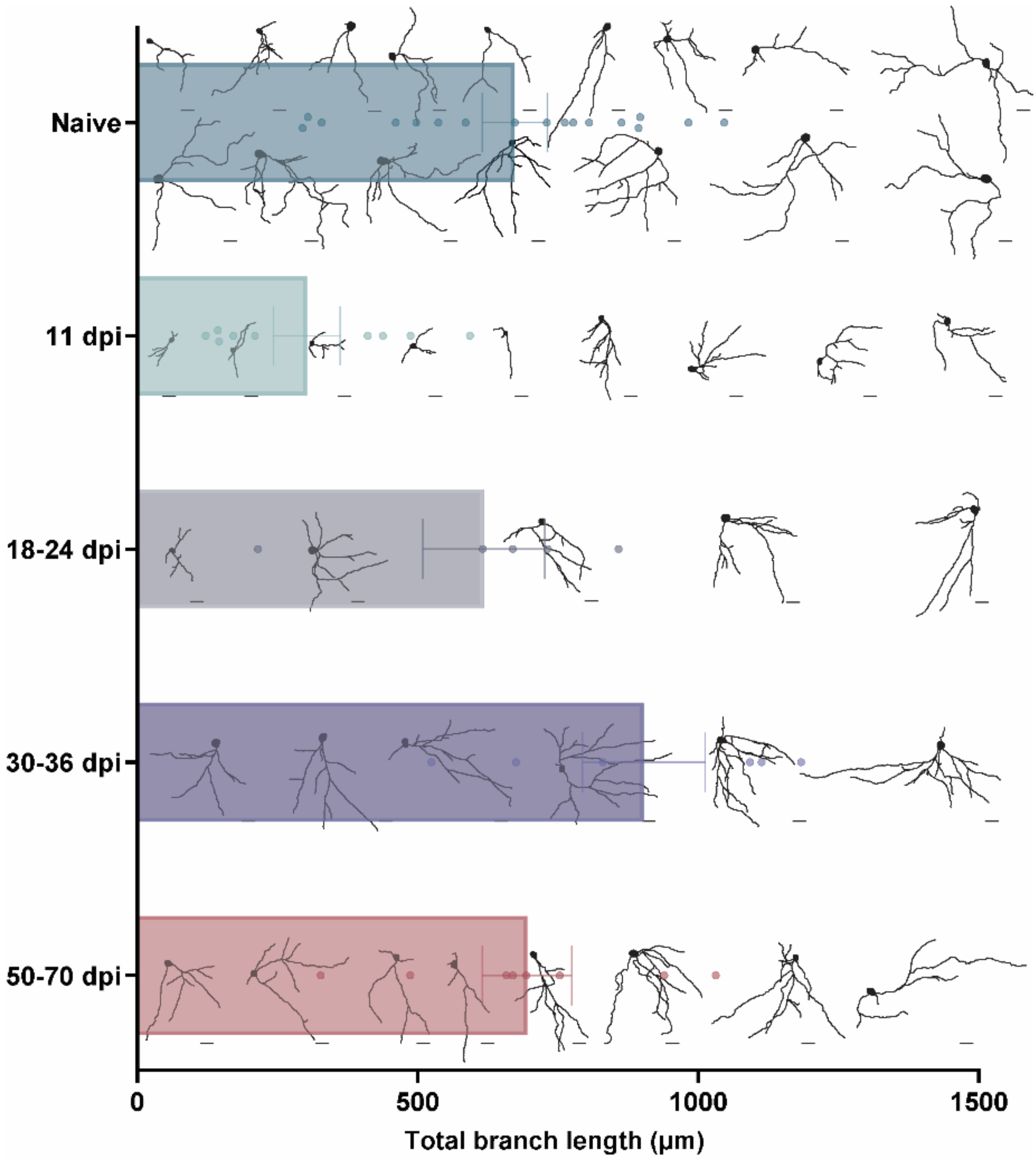


**Supplementary Figure 5: Total branch length of neurons determined using Sholl analysis and all skeletons of the traced cells per time point.** Naive neurons are transduced with an AAV 2.1 vector. All post-injury neurons are transduced with the gamma MLV retroviral vector. Scale bar: 20 µm

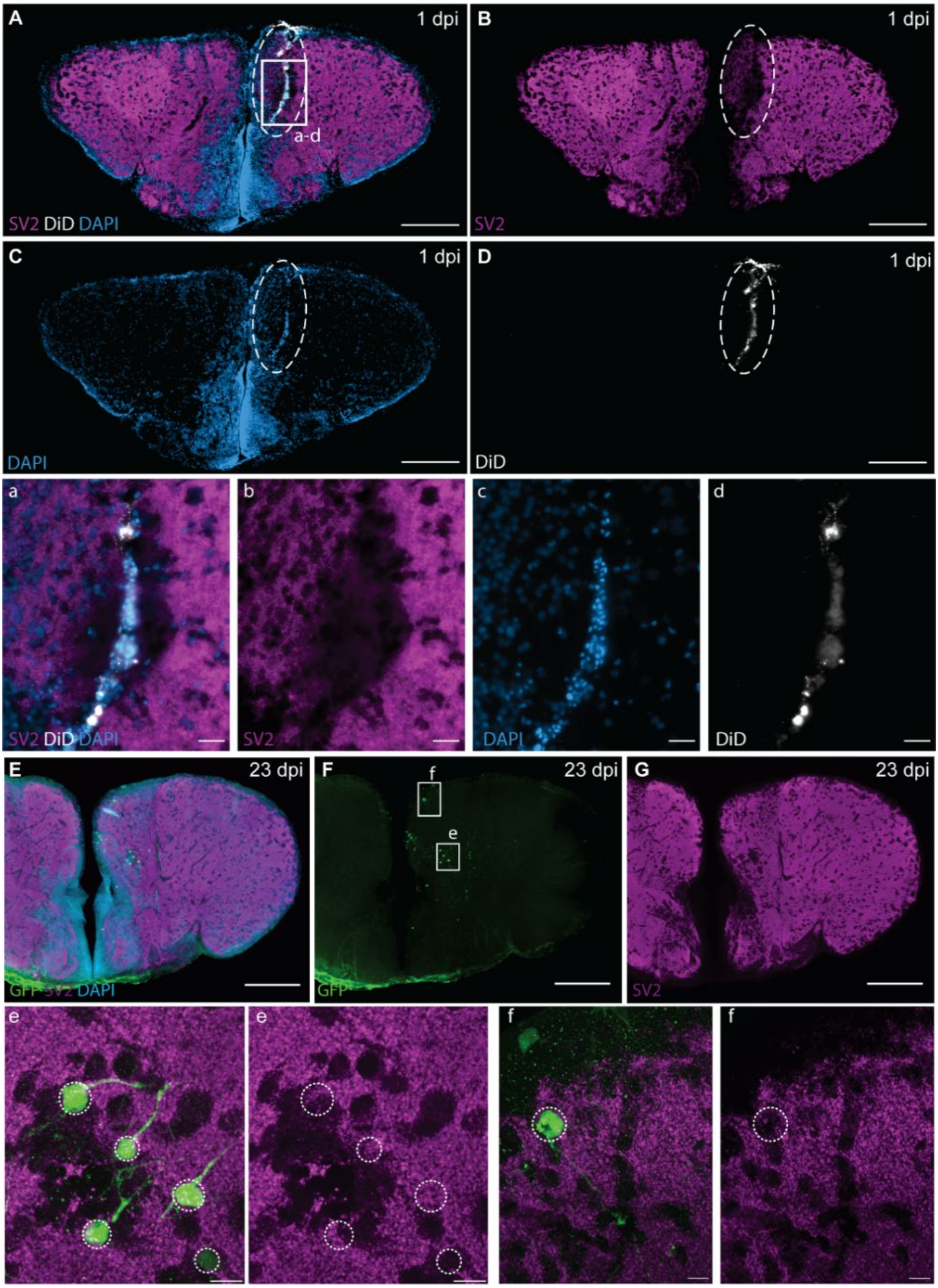


**Supplementary Figure 6: The number of SV2+ synaptic contacts is drastically reduced around the injury site at 1 dpi and reappears in normal amounts around the GFP+ newborn neurons by 23 dpi.** Coronal cryosections showing SV2 (synaptic vesicle 2 – magenta), DiD dye (white) and DAPI (blue) signal. At the core of the injury site, no synaptic contacts are detectable, and the part of the Dm zone left of the injury is also showing a clearly lighter signal intensity for SV2 (n=4). A-D: 20X Tile images,

Scale bar: 200 µm. a-d: 40X magnification, scale bar: 20 µm. (E-G) Fifty µm-thick vibratome sections showing SV2+ (magenta) punctae around the cell bodies and neurites of GFP+ newborn neurons by 23 dpi, indicative of a mature synaptic network involving the injury-induced newborn neurons. Scale bar: E-G: 200 µm; e,f: 10 µm.

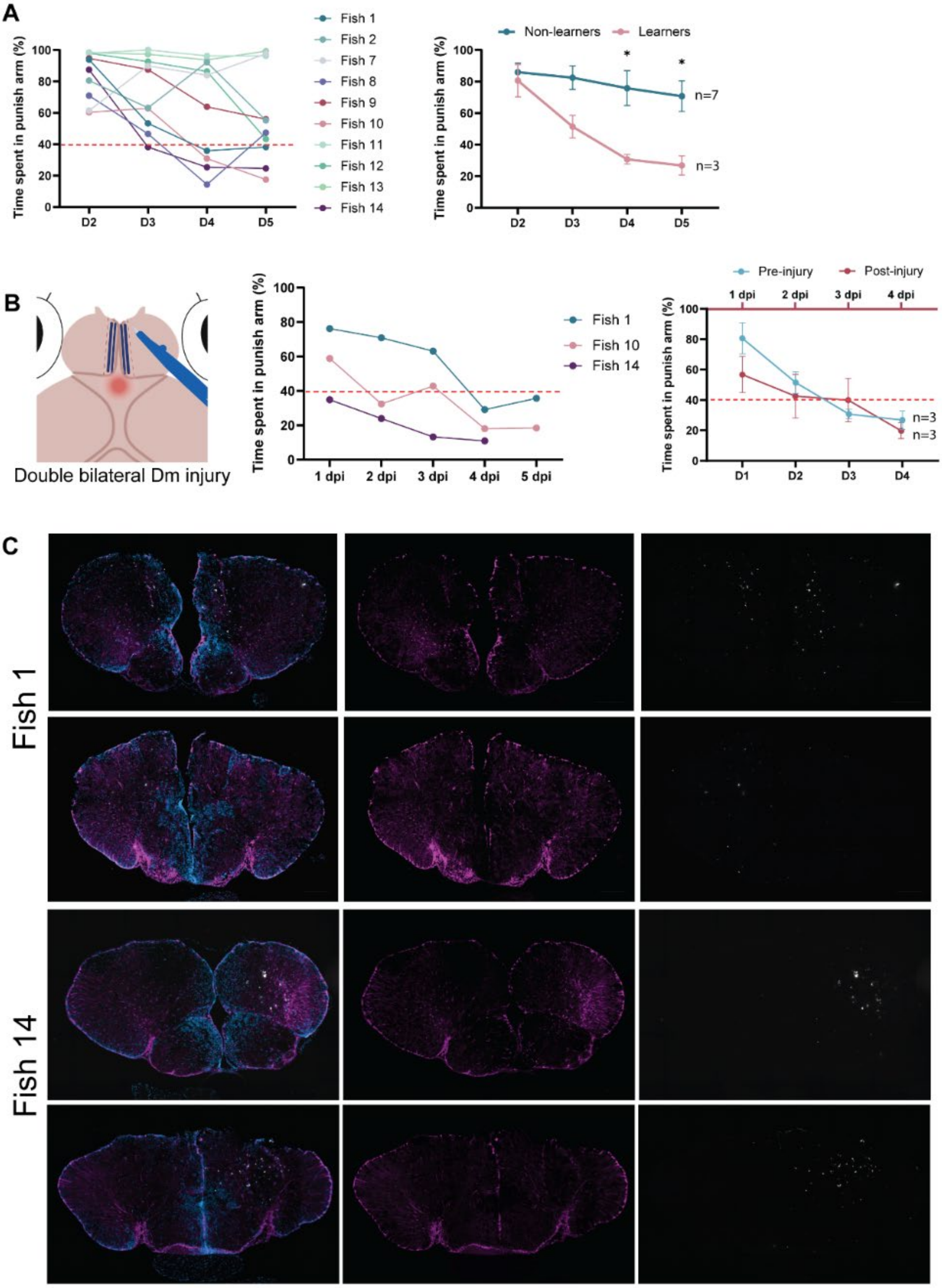


**Supplementary Figure 7: Injuries outside the Dm zone, unilateral lesions, or lesions that do not run along the complete anterior-posterior axis of the telencephalon do not impair memory in the first**

**days post injury.** (A) Using Learning paradigm 1 after dark preference pre-selection, 3 out of 10 fish of a new batch learned to avoid the dark arm. A significant difference between learners and non-learners was observed from the third testing day ($p_{D4}=0.0150$, $p_{D5}=0.0184$, two-way ANOVA). (B) Next the 3 learners each received a different lesion: a bilateral injury but either only the anterior zone of the Dm (Fish 1), or lateral from the Dm (Fish 10) or a correct albeit unilateral (right) Dm injury along the full anterior-posterior axis (Fish 14). All lesions have been confirmed by immunohistology for GS and DAPI. All 3 fish were still able to perform equally well in the conditioned place avoidance test, like in their learning week, in the week after lesioning, indicative of memory and learning retention, by relying on partially or fully functioning Dm zones.

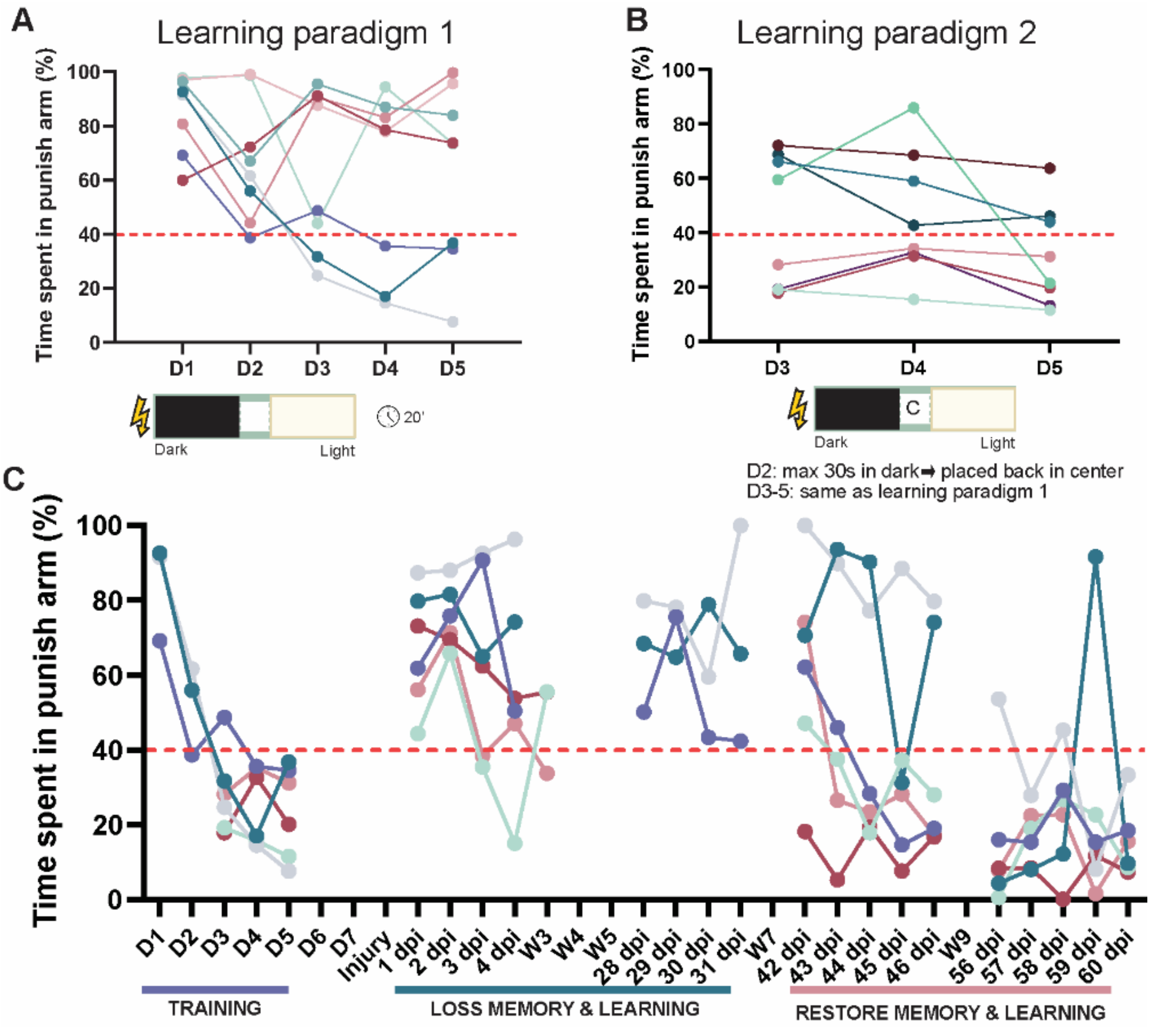


**Supplementary Figure 8: A different learning paradigm (nr. 2) did not lead to an increase in the number of learners but confirmed the role of the Dm in conditioned place avoidance learning.** (A) In learning paradigm 1, fish were allowed to swim freely for 20 min in the light-dark maze which is coupled to an electric shock if fish enter the dark arm from day 1. (B) In learning paradigm 2, fish were selected on day 1 on their dark preference and received one electric shock upon entering the dark arm. The 8 fish with the lowest latency to reach the dark were selected and continued with. On day 2, these fish were placed in the maze coupled to the electric shock in the dark zone. If fish stayed in the dark area for longer than 30s, they were placed back in the center zone 'C', for maximally 15 times. From day 3 onwards the procedure was identical to the one of learning paradigm 1. (C) Time spent in the punish/dark arm over the two different experiments and phases: training (day 1-5, n=6); injury on day 8; injury leads to loss of memory 1-4 dpi (n=6) up to 30 dpi (dpi, n=3); functional recovery via learning and memory is observed from 42 dpi up to 60 dpi (n=6).

**Supplementary Table 1: Padlock probe sequence sets used for in situ sequencing.**

| Probe Name | Sequence |
|---|---|
| P2RY12_1 | TCAGGACCATGCTGTGTAGTGTACTCCCAAGTCCTTGCGTCTATTTAGTGGAGCCTCCTCTGTCGGTCCT |
| P2RY12_2 | CGTGTCCTTGACATCGTAGTGTACTCCCAAGTCCTTGCGTCTATTTAGTGGAGCCGAACACTGTGCAAAA |
| P2RY12_3 | TACAAGTCCTACTCCGTAGTGTACTCCCAAGTCCTTGCGTCTATTTAGTGGAGCCATCAGCAAGGAGCTG |
| P2RY12_4 | CATGACCCTGACCTTGTAGTGTACTCCCAAGTCCTTGCGTCTATTTAGTGGAGCCTGCTGCTGATCTCAT |
| P2RY12_5 | TTGACCGCTGCAGAAGTAGTGTACTCCCAAGTCCTTGCGTCTATTTAGTGGAGCCTTGGCCTCATCAGTA |
| NeuroD2_1 | AGATAAGCTCCTTTGGACTTATACGTCATGCGACCTGCGTCTATTTAGTGGAGCCCTGCTTGAGATGATC |
| NeuroD2_2 | AGGATGAGCACGAGCGACTTATACGTCATGCGACCTGCGTCTATTTAGTGGAGCCATGGCTGCAAGGTGA |
| NeuroD2_3 | TCCTGACTGAACAGTGACTTATACGTCATGCGACCTGCGTCTATTTAGTGGAGCCTCAACACCAGGAACT |
| NeuroD2_4 | ACGACAGGAACTATCGACTTATACGTCATGCGACCTGCGTCTATTTAGTGGAGCCAGGAGTCCTCAGACA |
| NeuroD2_5 | AAGCAGGCTTAGATGGACTTATACGTCATGCGACCTGCGTCTATTTAGTGGAGCCCTGGGAAACTTGTCC |
| GLUL_1 | ATGCTGGAGTCAACAGCGTTTAATGTGAGCTACGATGCGTCTATTTAGTGGAGCCACAGGGCTTGTCTGT |
| GLUL_2 | AGTAGCGTAACGACAGCGTTTAATGTGAGCTACGATGCGTCTATTTAGTGGAGCCCCGATACAGTTCTAG |
| GLUL_3 | TCACAGCGAGACATGGCGTTTAATGTGAGCTACGATGCGTCTATTTAGTGGAGCCCTATGTTAAGCAGCT |
| GLUL_4 | TACCCAGCTTGTTGAGCGTTTAATGTGAGCTACGATGCGTCTATTTAGTGGAGCCACTTCTCTGCAGTCT |
| GLUL_5 | GCCCTGTTGCTATGAGCGTTTAATGTGAGCTACGATGCGTCTATTTAGTGGAGCCATGGAGAAGGATGCT |
| GLUL_6 | GTCCTGCAGCTGCATGCGTTTAATGTGAGCTACGATGCGTCTATTTAGTGGAGCCCCTGTGCTGTGATTG |
| PCNA_1 | GGCCTTGAAGGATCTCGAGATTACAATTCACGAGCTGCGTCTATTTAGTGGAGCCGAAGAAGGTGTTGGA |
| PCNA_2 | ATGGACTCCTCTCACCGAGATTACAATTCACGAGCTGCGTCTATTTAGTGGAGCCATCTCGCTGCAGAGC |
| PCNA_3 | TGAAATGTGCAGGAACGAGATTACAATTCACGAGCTGCGTCTATTTAGTGGAGCCGTATGTCAAAGATCC |
| PCNA_4 | CAGAGCAGGAGTACACGAGATTACAATTCACGAGCTGCGTCTATTTAGTGGAGCCAACAGCTCGGTATTC |
| PCNA_5 | AATGAGCCAGTCCAACGAGATTACAATTCACGAGCTGCGTCTATTTAGTGGAGCCGTCACCATTGAAATG |
| PCNA_6 | CTTCACAAAGGCCACCGAGATTACAATTCACGAGCTGCGTCTATTTAGTGGAGCCGAACTACCTGAACTT |
| VIM_1 | GTGAGGTGGACACTCGCTACCACCTACGTGTATATTGCGTCTATTTAGTGGAGCCGCTGATGCTGCAGAG |
| VIM_2 | GACCAGCTCCTCATTGCTACCACCTACGTGTATATTGCGTCTATTTAGTGGAGCCGTAAGATGTCCTACA |
| VIM_3 | ATGCAGTCGCTCAACGCTACCACCTACGTGTATATTGCGTCTATTTAGTGGAGCCAACGAGAAGGCGCAG |
| VIM_4 | CTCCCAGAGGGAAGAGCTACCACCTACGTGTATATTGCGTCTATTTAGTGGAGCCGCTGCAGGATGAAAT |
| VIM_5 | CAGGATGAGATTAACGCTACCACCTACGTGTATATTGCGTCTATTTAGTGGAGCCAAGGTCGAGTCACTC |
| VIM_6 | CGCCAAGTTCAAGTCGCTACCACCTACGTGTATATTGCGTCTATTTAGTGGAGCCGCCAATGACTACAGG |
| MKI67_1 | TTCTCAACAGGAGCTTAGTGCAGATGACGTTAGTCTGCGTCTATTTAGTGGAGCCATCTGGAGAGGCTGA |
| MKI67_2 | TGAGGACTTGACTGGTAGTGCAGATGACGTTAGTCTGCGTCTATTTAGTGGAGCCGGACATGAAGATGGA |
| MKI67_3 | ACCGTGAAGAGTCAATAGTGCAGATGACGTTAGTCTGCGTCTATTTAGTGGAGCCCTTAGTGTAGCTTCG |
| MKI67_4 | AGCCACCAACTTACTTAGTGCAGATGACGTTAGTCTGCGTCTATTTAGTGGAGCCAGGAGCTCATGCAGG |
| MKI67_5 | AGTGAGCAGCTGGAGTAGTGCAGATGACGTTAGTCTGCGTCTATTTAGTGGAGCCGCAAGTGAAACCTGC |
| MKI67_6 | AAAGTACCAGCTCGTTAGTGCAGATGACGTTAGTCTGCGTCTATTTAGTGGAGCCAGCCTGAAGAGCAGA |
| HMGB2A_1 | CTAGAGGGAAGACCTTCTTAGGTACTTGCAGTGACTGCGTCTATTTAGTGGAGCCAGGACCCGAATAAGC |
| HMGB2A_2 | GAAGACCATGTCAGCTCTTAGGTACTTGCAGTGACTGCGTCTATTTAGTGGAGCCATGTTCCGAAAGATG |
| HMGB2A_3 | AGGTGCCAAGAAAGGTCTTAGGTACTTGCAGTGACTGCGTCTATTTAGTGGAGCCCTACGTCCCTCCTAA |
| HMGB2A_4 | TGCCAAGAAGCTTGGTCTTAGGTACTTGCAGTGACTGCGTCTATTTAGTGGAGCCCTCCATCGGAGATGT |
| HMGB2A_5 | CCATACGAGGCTAGGTCTTAGGTACTTGCAGTGACTGCGTCTATTTAGTGGAGCCTCCAAAGACAAGGCG |
| HMGB2A_6 | GAGCAAAAGGAGGTTTCTTAGGTACTTGCAGTGACTGCGTCTATTTAGTGGAGCCATGTTGCTGCCTACA |
| APOEB_1 | CCATGAAGGCTCTGATAGTTCATACGCGGTAGACATGCGTCTATTTAGTGGAGCCGTCTCCATCTATCAA |

| | |
|---|---|
| **APOEB_2** | GTCCTCTCAGCTCACTAGTTCATACGCGGTAGACATGCGTCTATTTAGTGGAGCCTGTGGACAACCTCAA |
| **APOEB_3** | GACCTGCAACTTCTGTAGTTCATACGCGGTAGACATGCGTCTATTTAGTGGAGCCAACCAGATGTCCCTA |
| **APOEB_4** | CATCAACACCTACACTAGTTCATACGCGGTAGACATGCGTCTATTTAGTGGAGCCCAATGTGAAGAACCG |
| **APOEB_5** | AGCACATCCATGGACTAGTTCATACGCGGTAGACATGCGTCTATTTAGTGGAGCCAGCCTGAAGGATCTG |
| **APOEB_6** | GTCAAGGAGACTGCCTAGTTCATACGCGGTAGACATGCGTCTATTTAGTGGAGCCGACATCATGGACAAG |
| **GAP43_1** | AGAGGAGCCCAAGGTGTTAGAGTACGCTAGTTGCATGCGTCTATTTAGTGGAGCCCTCTGAACCTCACCA |
| **GAP43_2** | ATAGTATGAGGAAGTGTTAGAGTACGCTAGTTGCATGCGTCTATTTAGTGGAGCCCTTCTGCTCAGACTA |
| **GAP43_3** | ATTTGAATGCATTGCGTTAGAGTACGCTAGTTGCATGCGTCTATTTAGTGGAGCCGGAATAGTGACACAG |
| **GAP43_4** | GTCAGTTGTATCTTGGTTAGAGTACGCTAGTTGCATGCGTCTATTTAGTGGAGCCTGACATATCCAGCTG |
| **GAP43_5** | GCCTCTGTCAGCTTTGTTAGAGTACGCTAGTTGCATGCGTCTATTTAGTGGAGCCAAAAGCCCTAAGAAG |
| **GAP43_6** | TGGAAACTCTCCCTTGTTAGAGTACGCTAGTTGCATGCGTCTATTTAGTGGAGCCGAACCAGATGCATGA |
